# Model-based detection of whole-genome duplications in a phylogeny

**DOI:** 10.1101/2020.01.24.917997

**Authors:** Arthur Zwaenepoel, Yves Van de Peer

## Abstract

Ancient whole-genome duplications (WGDs) leave signatures in comparative genomic data sets that can be harnessed to detect these events of presumed evolutionary importance. Current statistical approaches for the detection of ancient WGDs in a phylogenetic context have two main drawbacks. The first is that unwarranted restrictive assumptions on the ‘background’ gene duplication and loss rates make inferences unreliable in the face of model violations. The second is that most methods can only be used to examine a limited set of *a priori* selected WGD hypotheses; and cannot be used to discover WGDs in a phylogeny. In this study we develop an approach for WGD inference using gene count data that seeks to overcome both issues. We employ a phylogenetic birth-death model that includes WGD in a flexible hierarchical Bayesian approach, and use reversible-jump MCMC to perform Bayesian inference of branch-specific duplication, loss and WGD retention rates accross the space of WGD configurations. We evaluate the proposed method using simulations, apply it to data sets from flowering plants and discuss the statistical intricacies of model-based WGD inference.

## Introduction

“Every gene from a pre-existing gene”, proclaimed Muller, echoing Virchow’s famous third dictum (Muller 1936). This principle, though no longer thought of as a rule without exception (Long et al. 2013; Ruiz-Orera et al. 2018; Zhang et al. 2019), is still the paradigm for thinking about gene content evolution; with gene duplication regarded as the primary driver. A diverse array of molecular processes cause gene duplication (and loss) (see for instance Lynch (2007)), and it is generally insightful to distinguish two main classes. Small-scale duplication and loss (SSDL) events, resulting in duplication or loss of a single or couple of genes, are thought to originate from processes such as non-homologous recombination and transposition. These may be contrasted with large-scale duplication and loss events involving the whole genome or a large fraction of it, which generally result from chromosome-level processes such as aneuploidy and rediploidization after polyploidization. We refer to the latter process by the colloquial term *whole-genome duplication* (WGD). It is nowadays well appreciated that ancestral WGD has been of considerable importance in the evolution of eukaryotic genomes (Van de Peer, Mizrachi, and Marchal 2017), and especially so in plants, where it would be safe to state that the genome of every extant plant has been shaped to some degree by ancient WGD events (1KP initiative 2019). Identifying ancient WGDs through comparative genomic analyses remains however a non-trivial task, as exemplified by several recent controversies (Li et al. 2019; Nakatani and McLysaght 2019; Zwaenepoel et al. 2019), illustrating the need for reliable statistical methods to detect WGDs in a phylogenetic context.

One of the consequences of Muller’s adage is that a particular class of stochastic models, namely birth-death processes (BDPs) (Feller 2015), have since long been a mainstay for modeling the evolution of gene family sizes (Novozhilov, Karev, and Koonin 2006). In particular the linear BDP and its variants have been widely used because of their tractable transition probabilities, facilitating statistical inference in a phylogenetic context (Hahn et al. 2005; Csűrös and Miklós 2009; Rabier, Ta, and Ané 2013; Liu et al. 2011; Librado, Vieira, and Rozas 2011; Tiley, Ane, and Burleigh 2016; Tasdighian et al. 2017). In these approaches, a series of linear BDPs are assumed to operate along the branches of a known species tree, providing an intuitive generative model for gene counts in a set of taxa for a single gene family. Such a set of gene counts is often referred to as a *phylogenetic profile* and can be obtained from genomic data sets using standard comparative genomics methods, thereby providing an attractive data set for likelihood-based inference of phylogenetic BDP models. As a model of gene family evolution, the birth and death rate parameters of the linear BDP are generally interpreted as the per-gene rate of duplication and loss per unit of time, thereby constituting a reasonable model of gene family evolution by SSDL (Novozhilov, Karev, and Koonin 2006). Interestingly, large scale duplication and loss, and in particular WGD, cause a characteristic deviation from the SSDL-driven evolutionary process. Indeed, posterior predictive simulations (performed using methods and data discussed below) clearly show that WGDs that are not accounted for are the main source of lack of fit of the phylogenetic linear BDP in a plant data set with several well-known WGD events (Figure S1, S2, S3). Furthermore, these unmodeled WGDs cause biases in the estimated duplication and loss rates, compromising their interpretation as rates of the SSDL process.

Rabier, Ta, and Ané (2013) were the first to propose a model of gene family evolution including ancestral WGDs, exploiting the deviation from a linear BDP to infer WGDs in a statistically founded approach. The same model was adopted in Zwaenepoel and Van de Peer (2019) in a gene tree reconciliation based inference context. In that study we showed that across-lineage variation in gene duplication and loss rates is a crucial factor that should be taken into account when performing model-based WGD-inference, and we developed a Bayesian approach modeling the variation in duplication and loss rates using relaxed molecular clock priors. In both Rabier, Ta, and Ané (2013) and Zwaenepoel and Van de Peer (2019), standard model selection techniques are used to determine whether some WGD hypothesis provides a significant better fit to the observed gene trees or phylogenetic profiles. These methods are therefore restricted to testing a limited set of *a priori* determined WGD hypotheses, and are unable to ‘automatically’ detect WGDs from the data. In this study, we develop an approach based on reversible-jump Markov chain Monte Carlo (rjMCMC) to infer WGDs under a probabilistic model of gene family evolution in a phylogenetic context based on a set of phylogenetic profiles without the need of specifying a restricted set of *a priori* WGD hypotheses. We study the performance of the new approach using simulations and explore its practical utility by applying the method to several comparative genomic data from plants.

## Methods

### Probabilistic model of gene content evolution

We employ a probabilistic model of gene family evolution based on the linear BDP to model the evolution of the number of genes in a gene family along a time-calibrated species tree 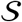 with 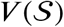 nodes (Hahn et al. 2005; Csűrös and Miklós 2009; Librado, Vieira, and Rozas 2011; Liu et al. 2011; Rabier, Ta, and Ané 2013; Tasdighian et al. 2017). Specifically, we assume an independent linear BDP for each branch *i* of 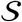 with duplication and loss rates *λ_i_* and *μ_i_*. We adopt the WGD model of Rabier, Ta, and Ané (2013) to introduce WGDs in the phylogenetic BDP, and call the resulting model the DLWGD model (for duplication, loss and WGD). Under the DLWGD model, we assume a set of *k* WGDs are indicated along 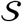, each characterized by a retention rate *q_j_* and age *t_j_*, *j* = 1,…, *k*. We denote by *b*(*j*) the branch of 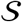 along which the WGD with index *j* is located. A set of *k* WGDs with the respective branches of 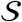 on which they are located will be called a ‘WGD configuration’. At the time of a WGD all extant gene lineages are duplicated, and a duplicated gene is either retained with probability *q_j_* or not retained with probability 1 − *q_j_*. Under this model, the complex polyploidization - rediploidization process associated with a WGD is assumed to happen in a time interval that is short relative to the branch length on which the WGD occurs, such that rediploidization is effectively instantaneous compared to the time-scale of the phylogeny and, crucially, has completed before the next speciation event. In other words, the number of genes retained in duplicate after rediploidization is modeled as a binomial random variable with ‘success’ probability *q_j_*. The resulting probabilistic graphical model (PGM) is a straightforward generative model for the evolution of gene trees (and as a consequence, also gene counts) evolving by means of duplication, loss and WGD with respect to an assumed species tree.

Parameter inference is based on a data set *X* consisting of *N* phylogenetic profiles, i.e. observed gene counts at the leaves of 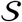. A single phylogenetic profile for family *i* is denoted as 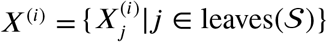, where we reserve subscripts for node indices and superscripts for family indices. We assume all *X*^(*i*)^, *i* = {1,…, *N*} are independent and identically distributed (iid) under the same DLWGD model. The linear BDP has a countably infinite state space, making direct computation of the likelihood under the DLWGD model using the pruning algorithm not possible. In most previous studies (Hahn et al. 2005; Librado, Vieira, and Rozas 2011; Tasdighian et al. 2017), this issue was circumvented by truncating the transition probability matrix to some carefully but arbitrarily chosen bound, where a trade-off exists between computational cost and accuracy. Alternatively, in a Bayesian approach one could augment the parameter space by the states at the internal nodes of 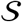 for every gene family, and sample from the augmented probability distribution using an MCMC algorithm, as was done by Liu et al. (2011). Following Rabier, Ta, and Ané (2013), we compute the likelihood for the resulting PGM using the algorithm of Csűrös and Miklós (2009), which uses *conditional survival likelihoods* in combination with a discrete prior distribution on the number of gene lineages at the root of a gene family to compute the marginal likelihood of a phylogenetic profile conditional on the assumed prior at the root. We employ a geometric prior on the number of lineages at the root with expected value 1/*η* following Rabier, Ta, and Ané (2013). For more details on the algorithm to compute the conditional survival likelihood under the DL and DLWGD model we refer the reader to Csűrös and Miklós (2009) and Rabier, Ta, and Ané (2013).

### Prior distributions and posterior inference

In previous work, we showed that modeling across-lineage variation in duplication and loss rates is crucial for the assessment of WGD hypotheses (Zwaenepoel and Van de Peer 2019). In that study, we modeled duplication and loss rates using a stochastic molecular clock by adopting priors similar to those used in Bayesian divergence time estimation. However, there we assumed the evolution of duplication and loss rates to be two independent processes. This assumption may be innocuous, but is not biologically plausible. Given that the expected value 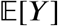 of the linear birth-death process over a time *t* for a given initial state *Y*_0_ is *Y*_0_*e*^(*λ*−*μ*)*t*^; for a gene family not to systematically expand or contract, *λ* should be approximately equal to *μ* for each branch in 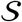 under the assumed model. A strong correlation between *λ* and *μ* is therefore expected *a priori* under what can be called a scenario of ‘neutral’ gene family evolution.

In the present work, we draw inspiration from Lartillot and Poujol (2010), and propose two bivariate models for across-lineage rate variation. The first model, faithful to Lartillot and Poujol (2010), models the evolution of *λ* and *μ* by a bivariate Brownian motion (BM) with some unknown covariance matrix Σ. The multivariate BM specifies a conditional probability density on the state of a random vector *θ_i_* = (log *λ_i_*, log *μ_i_*) for each *node i* of 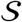, where we condition on the state of the parent node *j* at distance *t_i_*. Specifically, we have that

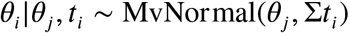

To obtain branch-specific duplication and loss rates, we approximate the sample path of the BM along a branch of the tree by taking the arithmetic average of the rates at the two flanking nodes. Alternatively, we consider a second model where rates across branches of 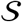 are independent and identically distributed, i.e.

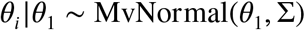

Where we denote the branch leading to node *i* with the index *i*, and denote for a branch with index *i* by *θ_i_* the vector of the logarithm of the branch-specific duplication and loss rates. In contrast to the bivariate BM model, this bivariate independent rates (IR) model can be specified directly in terms of branch-specific rates, resulting in considerable improvements in the efficiency of the MCMC algorithms we develop (see further). For both the BM and IR model we assume an inverse Wishart prior on the covariance matrix Σ with prior covariance matrix Ψ and degrees of freedom *q* = 3 (Lartillot and Poujol 2010). For the state at the root in the BM model, or the mean rates in the IR model (*θ*_1_), we assume a Multivariate Normal prior with covariance matrix Σ_0_ (which may or may not be chosen identical to Ψ) and prior mean vector *θ*_0_.

We denote the total tree length by *T* and assume a mapping from points in the interval [0, *T*] to points along 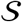, similar to for instance Huelsenbeck, Larget, and Swofford (2000). We assume a uniform prior on the interval [0, *T*] for the WGD times and a Beta prior for the associated retention rates *q*. Note that we treat the WGD times as random variables, whereas in previous studies these were usually assumed fixed, a potentially problematic assumption given the difficulties associated with dating uncertain WGD events (Rabier, Ta, and Ané 2013; Tiley, Ane, and Burleigh 2016; Zwaenepoel and Van de Peer 2019). Lastly, we consider a Beta prior on the *η* parameter that specifies the geometric prior on the number of lineages at the root. The full hierarchical model can be summarized as (taking the IR model as an example, see also Figure S4)

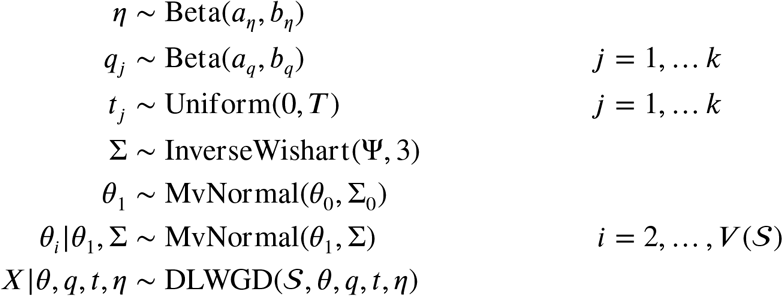

Unless stated otherwise, we used the following ‘vague’ prior settings: a Beta(3, 1) prior for *η*, a Beta(1, 3) prior for *q*, *θ*_0_ = [log(1.5), log(1.5)], Σ_0_ = [1 0.9; 0.9 1] and Ψ = *I*_2_, i.e. the 2 × 2 identity matrix. We make the important remark that any particular choice of prior parameterization for *θ*_0_, Σ_0_ and Ψ should depend on the time scale of the phylogeny. We sometimes consider an additional prior on the expected number of lineages at node *i* given one lineage at the parent node *j*, 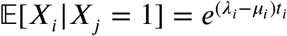, which serves to constrain the duplication and loss rate to a regime where they are of similar magnitude. For this ‘constraining’ prior we consider a Normal distribution with mean one and standard deviation *δ*. The advantage of this additional prior, which is not in a direct relation to the generative model, is that it allows to constrain duplication and loss rates in a biologically meaningful and insightful way due to its interpretation in terms of gene family expansion and contraction. Encoding such prior intuitions on gene family evolutionary dynamics in terms of the prior covariance matrix Ψ is generally less obvious.

We devised a Metropolis-within-Gibbs MCMC algorithm to obtain approximate samples from the posterior distribution for a model with a fixed WGD configuration. A single iteration of the MCMC algorithm involves a postorder traversal along the tree updating parameters using the following conditional updates:

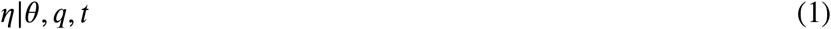

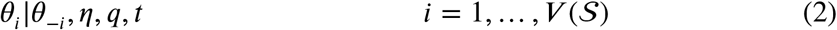

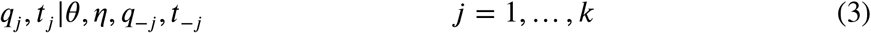

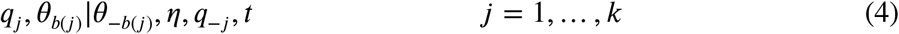

Where we use a common notational convenience by denoting by *θ*_−*i*_ the vector *θ* with *θ_i_* excluded. Note that we do not explicitly update the covariance matrix Σ, but use the fact that the inverse Wishart distribution is a conjugate prior for the multivariate normal to integrate out the covariance matrix Σ and compute 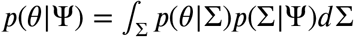 directly, as in Lartillot and Poujol (2010). For multivariate updates (2) and (4) we use a proposal strategy akin to Lartillot and Poujol (2010), where we propose a new state by executing one of the following moves uniformly at random: (a) additively update a uniformly chosen parameter using a uniform random walk proposal, (b) update all parameters by the same additive uniform random variable and (c) update all parameters additively with iid uniform random variables. For update (3), we update either *q_j_*, *t_j_* or both each with probability 1/3 using a uniform random walk proposal for *q_j_* and an independent proposal for *t_j_*, sampled from the time interval between the parent and child node of WGD *j*. Lastly, we update *η* using a uniform random walk proposal. For both *q* and *η* updates we reflect proposals that are out of bounds back into the (0, 1) interval. All proposal parameters were automatically tuned using diminishing adaptation during burn-in (Roberts and Rosenthal 2009).

### Reversible-jump MCMC

Previous work on the DLWGD model has considered the WGD hypotheses fixed and pre-specified, and resorted to standard model selection approaches using likelihood ratio tests or Bayes factors to select among a small set of nested candidate models or ‘WGD configurations’ (Rabier, Ta, and Ané 2013; Zwaenepoel and Van de Peer 2019). Here we treat the WGD configuration as unknown, and seek to jointly perform inference of branch-wise duplication and loss rates, the number of WGDs *k* and their locations and retention rates {*t_j_*, *q_j_*; *j* ∈ 1,…, *k*} from a set of exchangeable phylogenetic profiles 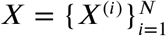 from *N* gene families.

Different configurations of WGD hypotheses along a phylogeny constitute different instances of the DLWGD model with the dimensionality of the parameter space depending on the number of WGDs on the branches of 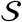. Denote by *ϕ_k_* the parameters associated with the DLWGD model with branch-wise duplication and loss rates *θ* and *k* WGDs. By Bayes’ theorem we have

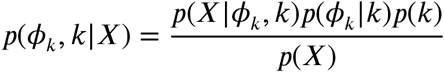

The reversible-jump MCMC algorithm (Green 1995) allows to simulate a Markov chain with *p*(*ϕ_k_*, *k*|*X*) as stationary distribution without requiring evaluation of the marginal likelihood *p*(*X*). In the rjMCMC algorithm, we construct trans-dimensional moves that add and remove WGDs to the current state. In the formalism of Green (1995), we denote the current state of dimensionality *n* as *ϕ*, and propose a new state *ϕ*′ of dimensionality *n*′ = *n* + 2 (i.e. a forward move, adding a WGD) by drawing a vector of random numbers *u* = (*u*_1_, *u*_2_, *u*_3_) from a joint density *g* such that (*ϕ*′, *u*′) = *h*(*ϕ*, *u*), where *h* is some deterministic function and *u*′ are the random numbers from a joint density *g*′ required for the reverse move from *ϕ*′ to *ϕ* with the inverse function *h*′ of *h*. Assume we introduce a WGD *j* on branch *b*(*j*) = *i*, we write *ϕ* = (*ϕ*\*, log *λ_i_*) and construct our forward move such that 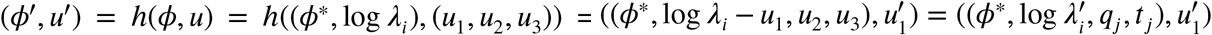. The acceptance probability for the forward move required for obtaining detailed balance is (Green 1995)

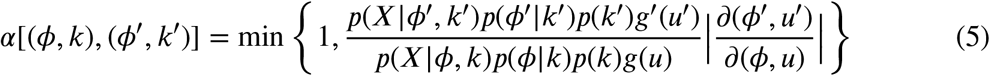

where, in our case, the absolute value of the Jacobian determinant is equal to one. We generate *u*_1_, *u*_2_ and *u*_3_ independently, so that the proposal density factorizes as *g*(*u*) = *g*_1_(*u*_1_)*g*_2_(*u*_2_)*g*(*u*_3_). The reverse move requires only one random variable 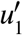, and we take 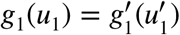. We further sample *u*_3_, i.e. the WGD location along 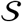, from the Uniform(0, *T*) prior. Collecting terms that appear in both the numerator and denominator and performing the appropriate cancellations, we obtain the acceptance probability of the forward move as

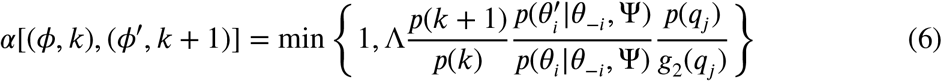

Where Λ denotes the likelihood ratio. Since we have a natural centering point in the sense of Brooks, Giudici, and Roberts (2003), i.e. Λ = 1 if *u* = *u*\* = **0**, we can see that under so-called weak non-identifiability centering, an optimal choice for *g*_2_(*u*_2_) may be to take the prior density *p*(*q*).

We further implemented two slightly different reversible jump kernels, (1) a kernel where not only *λ_i_* but also *μ_i_* is updated in a model jump and (2) a kernel where only a new WGD is proposed, without updating either *λ_i_* or *μ_i_*. The acceptance probability is identical to (6), and similar observations with regard to the choice of the proposal density of *q_j_* hold. By default we use the former of these two in our analyses. We verified the implementation of the MCMC algorithm by running the sampler in the absence of data, in which case the sample should approximately reproduce the prior (Figure S5, S6). Furthermore, we assessed our ability to recover accurate parameter estimates for data sets simulated under the DL and DLWGD models (see further).

### Posterior inference of WGD configurations

In a Bayesian framework, model comparisons are usually performed using Bayes factors. The Bayes factor of a model *M*_1_ vs. a model *M*_0_ is computed as

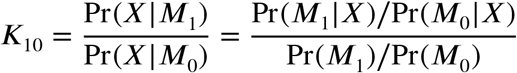

This is straightforward to compute given an approximate sample of the posterior distribution across model space. Note however that we have treated WGD configurations with the same number of WGDs *k* (and as a result the same dimensionality of parameter space) as a single model *M_k_*, yet model selection between different *k* is of limited interest. We are actually more interested in assessing whether some particular branch *e* is likely to be associated with a number *k_e_* of WGDs. The approximate marginal posterior probability *p*(*k_e_*|*X*) of *k_e_* WGDs on branch *e* is again easily obtained from the MCMC sample. Since the prior probability of a WGD on a particular branch *e* is proportional to the length of that branch *t_e_*, the marginal prior probability *p*(*k_e_*) of *k_e_* WGDs on branch *e* under a Geometric(*p*) prior on the number of WGDs in 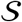 can be obtained as

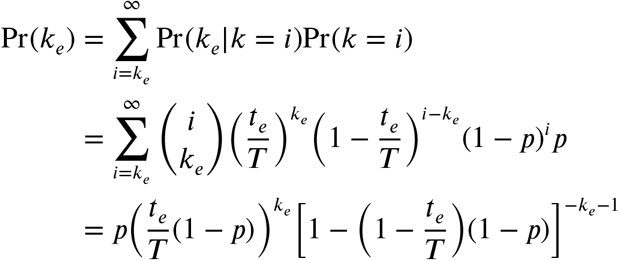

For a Poisson(*λ*) prior on *k*, a similar argument shows that 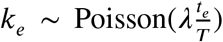, whereas for any discrete prior with finite support on *k*, the first equality gives a means to compute the relevant prior probabilities. This enables us to compute Bayes factors for branch-specific WGD configurations by comparing a WGD configuration with *k_e_* WGDs on branch *e* with a model with *k_e_* − 1 WGDs on that branch. This will be our main strategy for employing samples acquired using the rjMCMC algorithm for inference of WGDs along 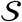.

### Posterior predictive model checks

We perform posterior predictive model checks by drawing 1000 random models and associated parameters from the approximated joint posterior distribution. For each such replicate we then simulate *N* phylogenetic profiles (i.e. a matrix with equal dimensions as the original data) from the associated DLWGD model. For these simulated data sets we compute a series of summary statistics (mean, standard deviation and entropy) for the numbers of genes at each leaf of 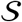 and the overall family size to acquire the posterior predictive distributions for these summary statistics. We then compare the observed values for these summary statistics with the relevant approximate posterior predictive distribution either graphically (e.g. Figure S17, S22) or numerically by computing posterior predictive *p*-values (Gelman et al. 2013).

### Gene family data

We obtained two data sets, one for a set of nine dicot species (*Arabidopsis thaliana, Carica papaya, Medicago truncatula, Populus trichocarpa, Vitis vinifera, Utricularia gibba, Chenopodium quinoa, Beta vulgaris* and *Solanum lycopersicum*) and another for a set of 12 monocot species (*Oryza sativa, Sorghum bicolor, Zea mays, Brachypodium dystachion, Ananas comosus, Elaeis guineensis, Musa acuminata, Asparagus officinalis, Phalaenopsis equestris, Apostasia shenzhenica, Spirodela polyrhiza* and *Zostera marina*). In this paper, in particular in figures, we refer to these species by three letter codes taking the first and first two letters from the genus and species names respectively (except for *Zostera marina*, where we take ‘zom’ to prevent collision with *Zea mays* (‘zma’)). All sequence data were gathered from PLAZA dicots 4.0 and PLAZA Monocots 4.5 (Van Bel et al. 2017). The dated species tree for the dicots was based on a tree with median node ages from TimeTree (Kumar et al. 2017), whereas for the monocots, divergence times were gathered from Foster et al. (2016). Gene families were then obtained using OrthoFinder with default settings. We filtered out families that did not contain at least one gene in each clade stemming from the root of the associated species tree, to rule out *de novo* origination of gene families in arbitrary clades of the species tree. We condition all our analysis on this filtering procedure as in Rabier, Ta, and Ané (2013) and Zwaenepoel and Van de Peer (2019). We further filtered out large families using a Poisson outlier criterion, filtering out families for which 2*Y* > median(2*Y*) + 3 where *Y* is the square root transformed family size. Unless stated otherwise, we restrict all our analyses to a random sample of 1000 gene families from the filtered data sets for the sake of computational tractability.

### Availability and implementation

All methods were implemented in the Julia programming language (v1.3, Bezanson et al. (2017)). Important computationally intensive routines support distributed computing in order to harness modern parallel computing environments. The associated software package is open source, documented, and freely available at https://github.com/arzwa/Beluga.jl. The data sets considered in this study are also available from that repository.

## Results

### Simulated data

We conducted simulation studies to evaluate the statistical performance of the new methods we propose. All simulations were performed for a species tree of nine eudicot species. We first verified our ability to recover true parameter values for data sets of 500 gene families simulated from the IR prior under a reasonable yet broad range of parameter values. For this first set of simulations, we randomly sampled covariance matrices from the inverse Wishart prior with Ψ = [0.5 0.45; 0.45 0.5] for 20 replicates. We then sampled branch-wise duplication and loss rates, assuming a multivariate lognormal prior with mean 1.5 and the same prior covariance matrix on the mean duplication and loss rates. These prior settings were based on preliminary analyses of a data set for the same species tree. For every replicate we sampled a random set of WGDs with uniform retention rates, using a Geometric prior with *p* = 0.2 on the number of WGDs in the phylogeny. Values for *η* were sampled from a Beta(5,1) prior. We then performed posterior inference using fixed-dimensional MCMC assuming the simulated WGD configuration, with Ψ = [1 0.5; 0.5 1], a prior mean duplication and loss rate of 1.5 and Σ_0_ = Ψ. For each replicate we sampled for 11000 iterations, discarding 1000 iterations as burn-in. Results for these simulations are shown in Figure 1. We find that overall, even in the presence of considerable across-lineage rate variation, branch-wise duplication and loss rates can be estimated with fair accuracy and with no obvious biases across approximately three orders of magnitude. Retention rates are similarly well estimated for all but a handful of WGDs. In cases where multiple WGDs were present on the same branch, estimated retention rates were sometimes swapped (e.g. the inferred retention rate for the first simulated WGD corresponded to the retention rate for the second WGD on the same branch and vice versa), explaining the handful of seemingly incorrectly estimated retention rates (Figure S7).

**Figure 1:**
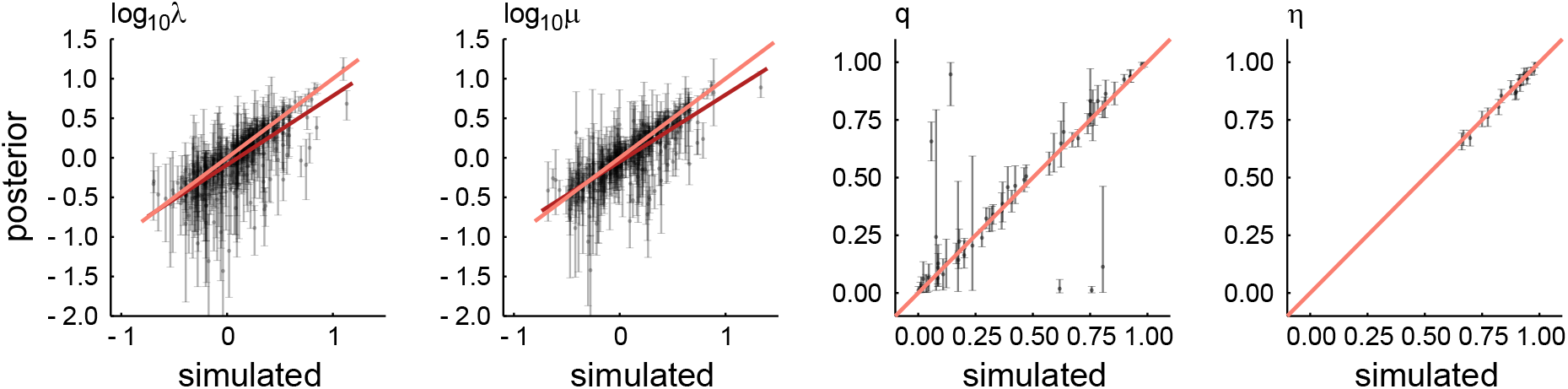
Comparison of posterior means and 95% credibility interval with simulated values. Results across different simulated data sets are pooled together in one plot. In orange, the ideal (expected) relation between posterior means and true (simulated) parameter values is shown, while in red the least squares regression of the posterior means on the true values is shown. We refer the reader to Figure S7 for plots of the retention rate estimates by replicate.

Next we studied the degree to which the assumption of constant rates across gene families affects our posterior inferences. For these simulations we simulated for each replicate genomewide lineage specific rates *θ* from a multivariate normal distribution with a fixed covariance matrix with variance 0.1 and covariance 0.09. This allows for limited, though not negligible, rate heterogeneity across lineages. For each replicate, we sampled a series of 500 relative rates (*r*) from a Gamma(*α*, 1/*α*) distribution with mean 1 and parameter *α*, with log_10_ *α* ∈ {−1,0. 5,0, 0.5, 1} multiplying for every gene family *i* the genome-wide rates by *r_i_*. For every replicate we thus obtained a set of 500 phylogenetic profiles from these family-specific BDPs. We then performed posterior inference under the IR model identical to our approach for the first set of simulations. These simulations indicated that for strongly asymmetric distributions of family-wise relative rates (small values of *α*), the genome-wide rates were systematically underestimated (Figure S8). If the distribution of relative rates is not too extreme however, the genome-wide average lineagespecific duplication and loss rates were well recovered even though our inference is ignorant of across-family rate heterogeneity. It is relatively straightforward to include simple mixture models for across family rate variation, such as Yang’s discrete Gamma model (Yang 1994), however this entails a considerable increase in computational time required for computing the likelihood of the data. As our models restricted to lineage-specific variation perform well if the distribution of rates across families is not too asymmetric, we do not further consider modeling family variation in this study.

To assess the performance of the reversible-jump MCMC algorithm for the inference of WGDs and branch-wise duplication and loss rates, we performed two sets of simulations with different branch-wise duplication and loss rates. For the first set of simulations, we simulated duplication and loss rates from the IR prior with a covariance matrix Ψ = Σ_0_ = [0.1 0.05; 0.05 0.1] and *θ*_0_ = [log(1.5) log(1.5)], again allowing for limited across-lineage rate heterogeneity. For the second set of simulations, we first obtained a sample from the posterior distribution for the dicot data set using a fixed DLWGD model with well-known WGDs marked along the branches of the species tree (*i.e. P. trichocarpa, C. quinoa, A. thaliana α* and *β*, *M. truncatula, S. lycopersicum* and three *U. gibba* WGDs) (Figure S9). For each simulation replicate, we then obtain a parameterized DLWGD model by drawing random branch-wise duplication and loss rates from the joint-posterior distribution acquired from the fixed-dimension MCMC sampler. Using this strategy, we ensure that realistic duplication and loss rates are used, allowing us to more properly assess the performance of the rjMCMC approach for WGD inference in reasonable settings, albeit tailored towards the dicot data set. In both sets of simulations, we randomly add six WGDs uniformly along the species tree with retention rates drawn from a Beta(1.5, 2) distribution. For each simulation scheme, we simulated 10 data sets of size *N* = 100, *N* = 500 and *N* = 1000 respectively to further assess the impact of using different amounts of data. We performed posterior inference using the default priors settings discussed in the methods section, with a discrete uniform prior on the number of WGDs ranging from 0 to 10 WGDs (note that this will limit the number of possible false positives). We performed three sets of inferences with the IR prior using different parameterizations of the prior covariance matrix with Ψ = *σ*^2^ *I*_2_ and *σ*^2^ ∈ {1, 0.2, 0.1}. We additionally perform inference using a constant-rates model where we assume the tree-wide duplication and loss rate are distributed as *θ*_1_ in the IR model. All results are based on 10000 iterations of the rjMCMC algorithm after discarding an initial 1000 samples as burn-in.

We observe fair performance, and as expected, reliability increases considerably when going from data sets of 100 to 500 families (Figure 2). The decrease in the false positive rate (FPR) is however considerably less, or even absent, when going from 500 to 1000 gene families, especially in the first set of simulations. Similarly, the power to detect a WGD does not seem to increase when using more data. Decreasing the variance of the bivariate process from *σ*^2^ = 1 to *σ*^2^ = 0.2 or *σ*^2^ = 0.1 does not seem to influence power nor FPR much for the first set of simulations (with limited across lineage rate heterogeneity), with the constant-rates model performing similar as the IR model under all settings for the prior covariance matrix. This is remarkable, as it suggests that using a prior that allows strong rate variation across lineages does not affect performance when these rates show in fact limited variation across the species tree. Prior assumptions on the rate evolution process do however affect performance for the second, arguably more realistic simulation set. Here we clearly observe the effect of assuming a restrictive model of rate variation across the tree, with the FPR increasing with decreasing *σ*. For the constant-rates model the FPR quickly rises to dramatic levels, and we note that here the prior on the number of WGDs in the species tree (a discrete uniform distribution from 0 to 10) prevents the number of false positive WGDs to rise even further. At the same time, the power is not considerably higher for more restrictive models on the rate evolution process, at least for reasonable Bayes factor thresholds.

**Figure 2:**
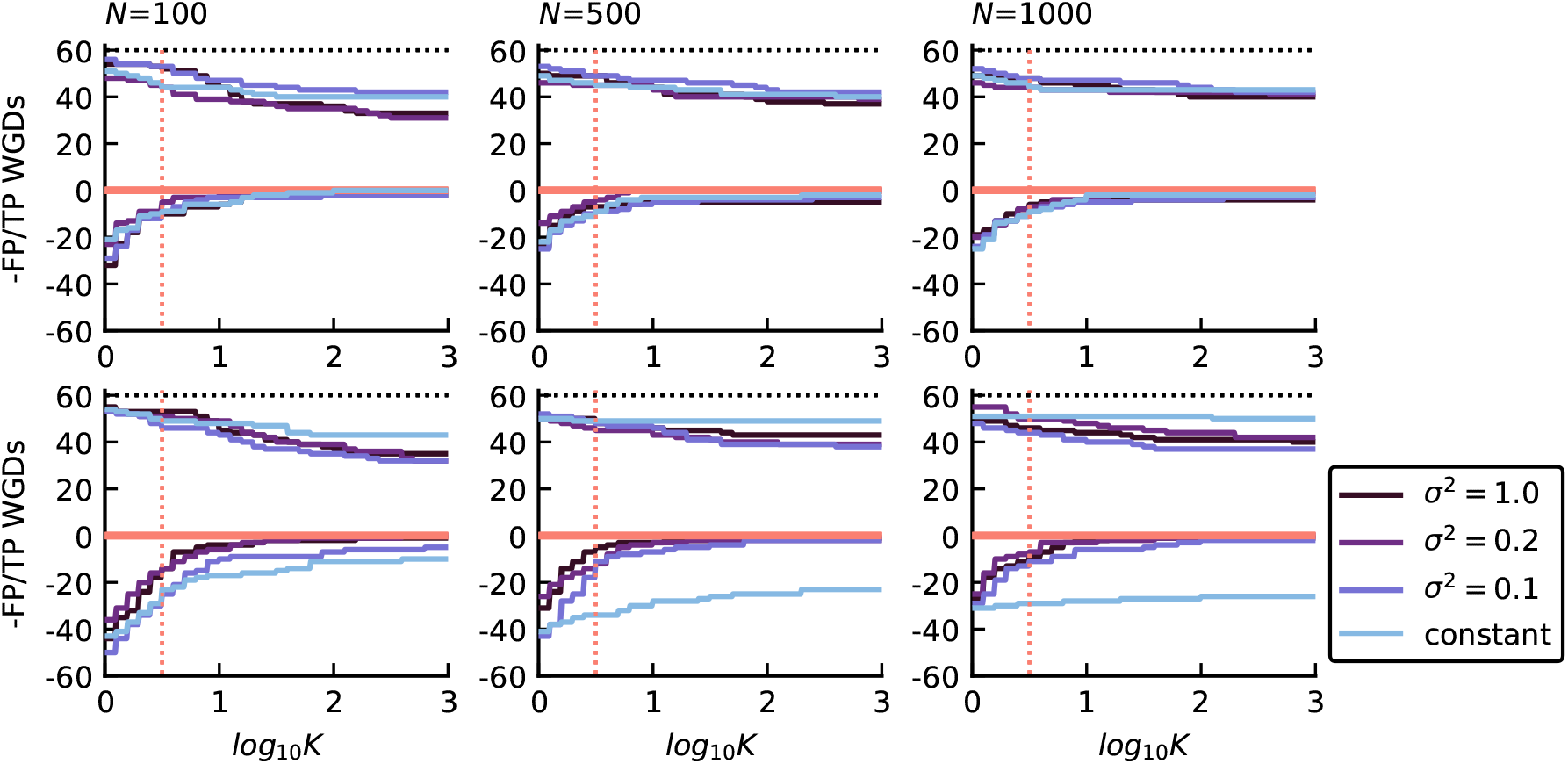
Performance of the rjMCMC sampler to detect simulated WGDs in different simulated data sets. The number of true positive (TP) and false positive (FP) WGDs detected for increasing cut-offs of the Bayes factor (log_10_ *K*) is shown by the positive and negative curves respectively. The top row shows the simulations from the IR prior, whereas the bottom row shows the simulations based on the joint posterior distribution for the dicots data set (see main text). In both series we show results for data sets of size *N* = 100, *N* = 500 and *N* = 1000 and a prior covariance matrix of *σ I*_2_, with *σ*^2^ ∈ {1, 0.2, 0.1}. We include results for a constant rates prior as well. Results are pooled across 10 replicate simulations. The black dotted line shows the total number of WGDs in the 10 replicates for each simulation. The red dotted line marks the usual rule-of-thumb threshold of 0.5 for the base 10 logarithm of the Bayes factor (log_10_ *K*).

A main challenge in rjMCMC is to obtain decent mixing across model space (Rannala and Yang 2013), and using more data may compromise our ability to jump between different WGD models. We record that for the simulations from the IR prior the acceptance probability of a between-model move was about 0.20, 0.09 and 0.06 for the *N* = 100, *N* = 500 and *N* = 1000 data sets respectively, illustrating this phenomenon. This could cause the power not to increase with increasing data set size, as would be expected, due to a decrease in the efficiency of the rjMCMC sampler to explore model space. We note that these simulations may exhibit such a phenomenon. Lastly we observe that overall, marginal posterior distributions for duplication and loss rates tend to agree with the simulated values, however for small data set sizes branch-wise duplication rates are considerably ‘shrinked’ towards the mean rate (Figure S10). Investigating some replicates more closely (e.g. Figure S11, S12), we make two observations: (1) duplication and loss rates for a branch are generally harder to obtain accurately in the presence of WGD(s) on that particular branch and (2) duplication and loss rates for short internal branches tend to be biased towards the tree-wide mean rates (*θ*_0_). We note that the latter observation reflects a feature that is not undesirable, because for short branches there will generally be less information in the data to accurately infer duplication and loss rates, and due to the hierarchical structure of our model these estimates tend to undergo ‘shrinkage’ towards the tree-wide mean.

### Plant data sets

We applied our new approach to two data sets from different plant taxa, focusing on well-sequenced genomes in dicots and monocots. These regions of the angiosperm phylogeny have been well-studied in terms of ancient WGDs, and, to some extent, a scientific consensus has emerged concerning which clades share particular ancestral WGD events, although considerable uncertainties remain (Ming et al. 2015; Jiao et al. 2014; Van de Peer, Mizrachi, and Marchal 2017; Soltis and Soltis 2016; Zhang et al. 2017). Based on some exploratory pilot runs, we use the IR prior with a prior covariance matrix Ψ = 0.5 *I*_2_, *θ*_0_ = [log 1.5, log 1.5], Σ_0_ = [1.0, 0.9; 0.9, 1.0], *q* ~ Beta(1, 3) and *η* ~ Beta(3, 1), with the additional constraining prior on the expected number of lineages per ancestral lineage using *δ* = 0.1 (unless stated otherwise). Throughout, we use a discrete uniform prior ranging from 0 to 20 for the number of WGDs. All results are based on 50000 iterations of the rjMCMC algorithm, of which 1000 were discarded as burn-in.

Applying the rjMCMC algorithm to the dicot data set (Figure S13, S19) revealed, as expected, strong support for WGD in poplar and quinoa, with in both cases posterior probabilities > 0.9 for the one-WGD model (Figure 3). These WGDs are associated with very high retention rates, with a marginal posterior mean and 95% credibility interval 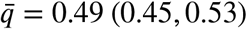 for the poplar WGD event, and 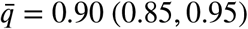 for the quinoa WGD. For *Arabidopsis*, we likewise find strong support for a WGD event, with the MAP model (at *P* ≈ 0.7) the one-WGD model and a marginal posterior mean retention rate at 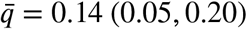. We do not find strong support for a second WGD on this branch, indicating that gene count data may not contain sufficient signal to detect multiple WGDs on a single branch if these are associated with relatively low retention rates. We observe that for most branches, there is very strong evidence *against* WGD with the Bayes factor strongly favoring the no-WGD model. Interestingly, the two branches for which the Bayes factor is about one (and thus indecisive both ways), namely the branches leading to *Solanum* and *Medicago*, are branches associated with WGD. The increased duplication rates for these branches suggest that the SSDL model is sufficiently flexible to accommodate the WGD-specific signal for long branches (Figure 4). Furthermore and perhaps unsurprisingly we find a strong bimodal posterior distribution for the duplication rate on both branches, with the modes associated with the competing no-WGD and one-WGD models (Figure S15). We further note that retention rates for the *Solanum* and *Medicago* events conditional on the one-WGD model for these branches were both estimated at 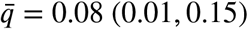, which is virtually identical to the results obtained with a fixed-dimensional MCMC sampler, suggesting that our inability to find decisive support for these WGDs is not due to issues in the rjMCMC algorithm.

**Figure 3:**
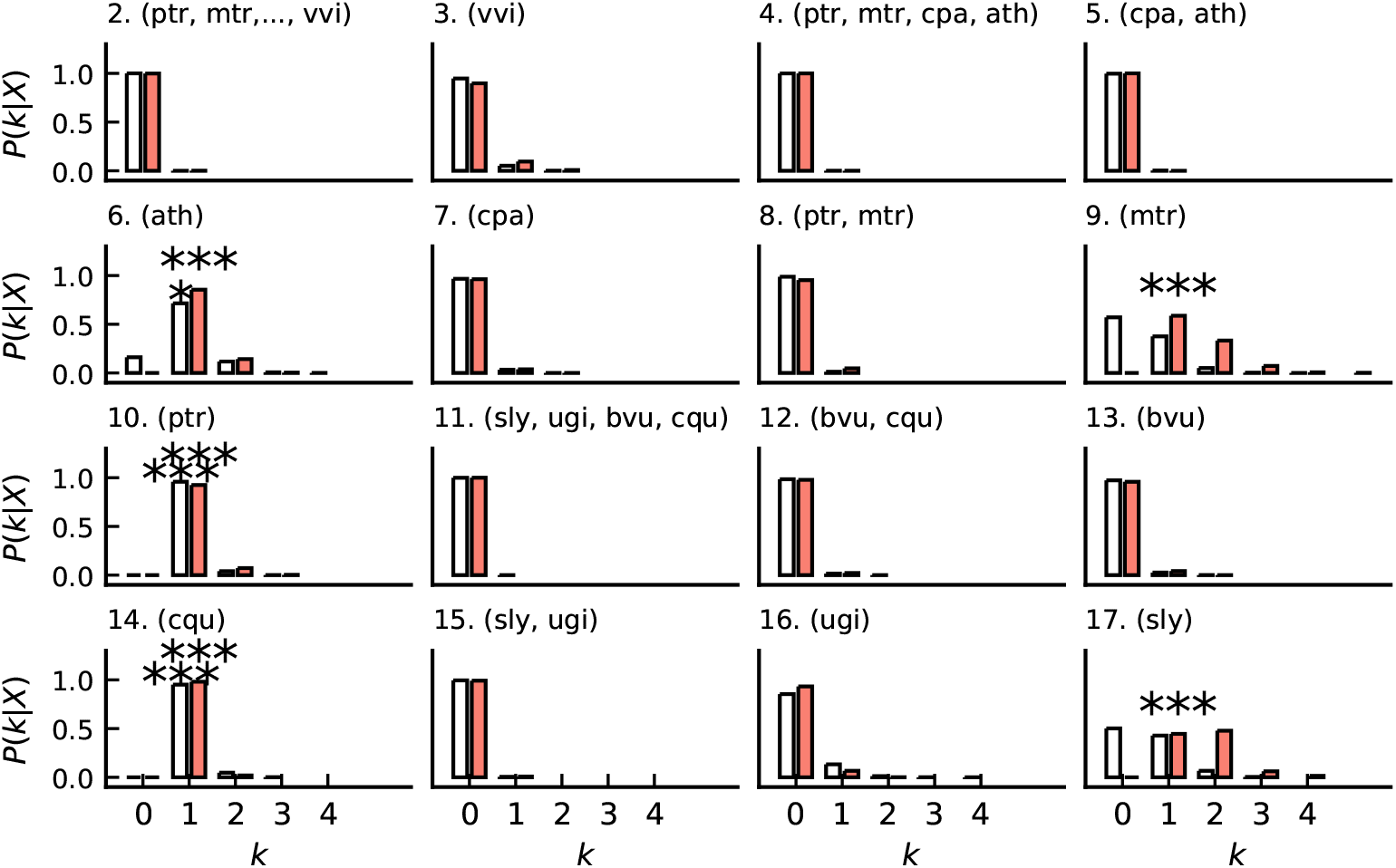
Posterior probability of the number of WGDs on each branch in the dicot data set for an analysis with *δ* = 0.1 (white) and an analysis with *δ* = 0.05 (red). Asterisks indicate the magnitude of the Bayes factor (*) 0.5 < log_10_ *K* < 1; (**) 1 < log_10_ *K* < 2; (***) log_10_ *K* > 2. Three letter codes (see methods) in between brackets denote the relevant clade below the particular branch.

**Figure 4:**
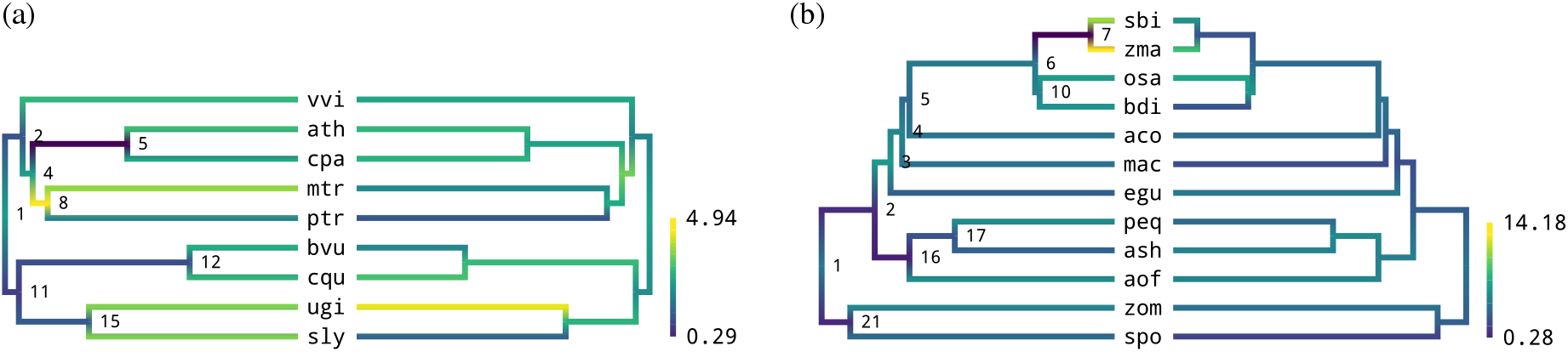
Marginal posterior mean duplication (left tree in each panel) and loss rates (right tree in each panel) on a scale of log_10_ (events per lineage per billion year) for (a) the dicot data set (*δ* = 0.1 analysis) and (b) the monocot data set (*δ* = 0.05 analysis). Estimates correspond to the across-model geometric average.

Assuming a stronger constraining prior on the expected number of lineages at the end of each branch, more specifically by setting *δ* = 0.05, resulted in very strong support for these two WGDs (Figure 3), with posterior retention rates of 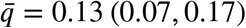 and 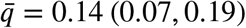 for the *Solanum* and *Medicago* WGDs respectively. Posterior inferences for other branches are almost identical to the *δ* = 0.1 analysis (Figure S13, S19), except for *A. thaliana*, where the posterior probability for the no-WGD model decreased to about zero (Figure 3), while the retention rate associated with the one-WGD model did not differ 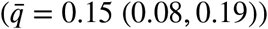. This analysis again clearly illustrates how phylogenetic WGD inferences are strongly dependent on prior assumptions on the SSDL process. Lastly, we note that we do not find support for any of the WGDs in *U. gibba* in any of our analyses, whereas the consensus view is that this lineage has undergone three WGD events since its divergence from other Lamiales (Ibarra-Laclette et al. 2013); a view that was largely established based on comparative co-linearity analyses. This lineage presents however a serious challenge to our approach, as its evolution has been associated with strong genomic reduction, characterized by, among others, extensive gene loss (Ibarra-Laclette et al. 2013; Carretero-Paulet et al. 2015). Our results suggest, in accord with this evolutionary history, that the purely quantitative signal from these WGDs has eroded considerably, and that they cannot be inferred from gene count data alone. We confirm the increased loss rate in this lineage, with an expected number of genes at the end of the branch per ancestral gene of 0.87 (0.80, 0.92) (Figure S16). Changing *δ* from 0.1 to 0.05 did not considerably alter this expected value, increasing it slightly to 0.89 (0.84, 0.93). Overall, posterior predictive simulations indicate a fairly good model fit, with 25 out of 30 of the observed summary statistics in the 95% central mass of the empirical posterior predictive distributions (Figure S17). In terms of the posterior predictive distributions there were no noticable differences between the chain with *δ* = 0.1 and *δ* = 0.05.

We performed the same analyses for the monocot data set (Figure 4, 5). We first performed the analysis with *δ* = 0.1 but were unable to recover the well-known WGD in the maize lineage and observed several branches for which the expected number of lineages per ancestral lineage under the SSDL process alone was relatively high due to high estimated duplication rates (mainly in the banana (*Musa*) and maize (*Zea*) lineages, Figure S21). An analysis with *δ* = 0.05 resulted in an inferred SSDL process that was more or less in equilibrium, and showed very strong support for the WGD in the maize lineage. We further report results for this second analysis with *δ* = 0.05. We find strong support for the so-called *τ* WGD event shared by all monocots in our data set except the Alismatales (branch 2 in Figure 4, 5), albeit with a fairly low retention rate 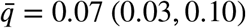. As in Zhang et al. (2017), but unlike some other studies (Ming et al. 2015; Jiao et al. 2014), we find support for a shared WGD event in the ancestor of all commelinids (branch 3), which we suggest to be the event referred to by *σ*, generally thought to be shared by all Poales (Ming et al. 2015). We note however that trace plots indicate that mixing across model space seems to be challenging for this branch (Figure S18), resulting in rather uncertain estimates for the posterior probability of this WGD. Again, this putative WGD event is associated with a relatively low retention rate 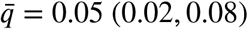. We find no support for the cereal-specific genome duplication event referred to by *ρ* (Figure 5, branch 6) in our gene count data, and find that the SSDL process along this branch is more or less in equilibrium, with the duplication rate 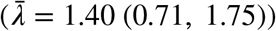 only slightly higher than the loss rate 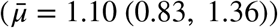 along this branch (Figure 4, S21). This is unlike our observations for the *Medicago* and *Solanum* branches in the nine-taxon dicot data set, where the absence of decisive support for the WGD events in these lineages was associated with increased duplication rates. The results for these putative ancestral WGD events in the monocot data set could indicate that the power of the gene count based rjMCMC approach for detection of ancestral WGDs might be affected by taxon-sampling issues, where WGDs on short branches leading to a species rich crown group can be detected more easily.

**Figure 5:**
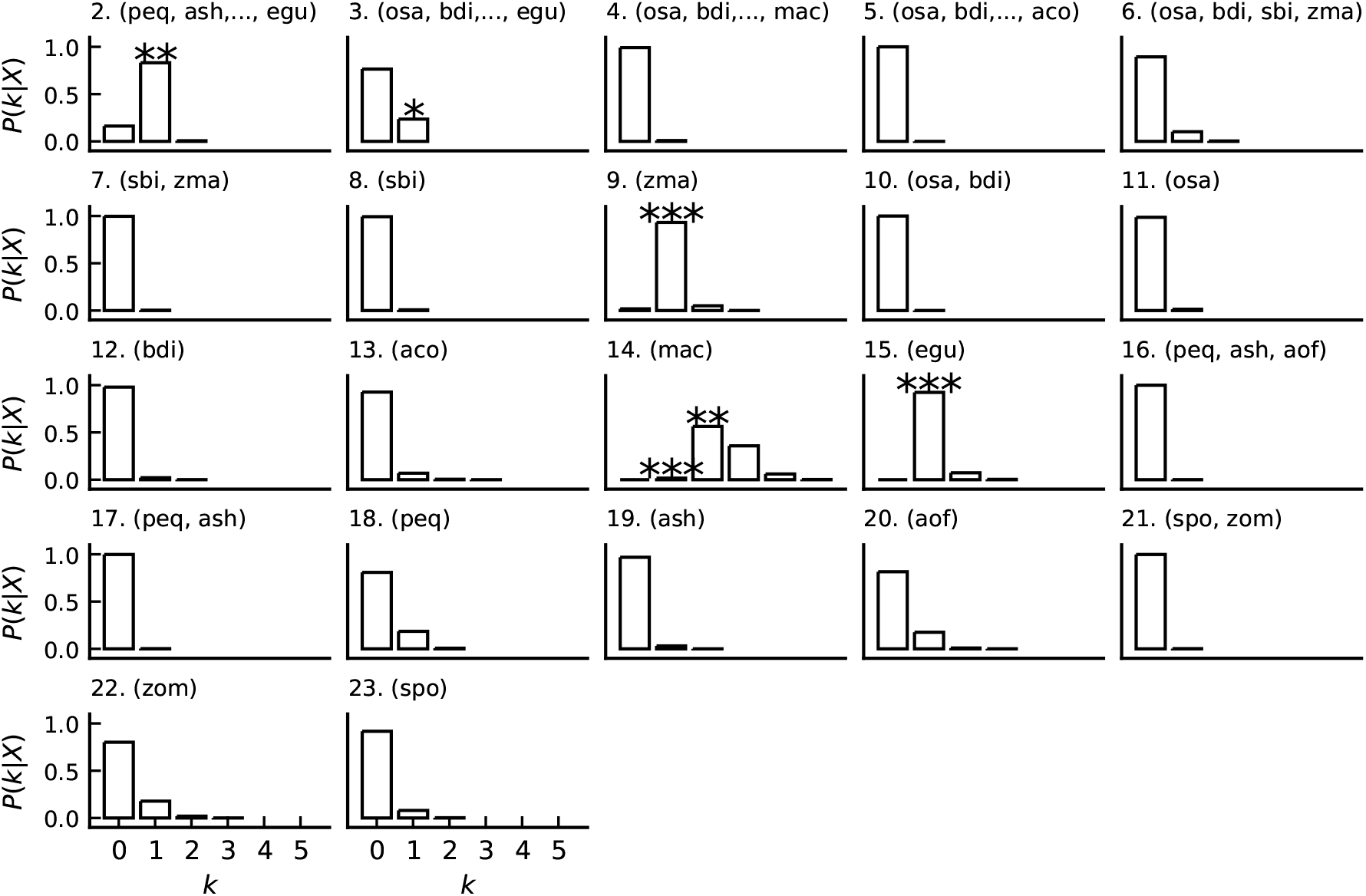
Posterior probability of the number of WGDs on each branch in the monocot data set (*δ* = 0.05 analysis). Interpretation is as in Figure 3. For species name abbreviations, we refer the reader to the methods section.

As already indicated, we find very strong support for WGD in the maize (*Zea*, 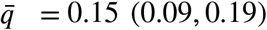) lineage when the prior with *δ* = 0.05 is used. The difficulty associated with identifying this WGD has likely to do with the high duplication rate in this lineage (Figure 4), which is notably higher than the rate in all other lineages in the monocot phylogeny considered here. We further find overwhelming support for WGD in the oil palm (*Elaeis*, 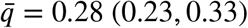) lineage, in line with previous results (Jiao et al. 2014; Singh et al. 2013). We also recovered the well-known multiple events along the branch leading to *Musa* (D’Hont et al. 2012), with strong support for a two-WGD model in the *δ* = 0.1 analysis and a three-WGD model in the analysis with *δ* = 0.05. The different WGDs can however not be distinguished, as exemplified by the fully overlapping distributions of the retention rates for the WGDs when ordered by the WGD time (Figure S20). Again, we find very strong evidence *against* WGD for most other branches, notably the branch leading to the orchids where a WGD was hypothesized by Zhang et al. (2017). The only branches that have some posterior probability for a one-WGD model (apart from the already discussed *ρ* WGD) are those leading to *Asparagus, Phalaenopsis, Zostera* and *Spirodela*. These lineages are thought to have underwent ancient WGDs (Olsen et al. 2016; Cai et al. 2015; Wang et al. 2014; Harkess et al. 2017), and are in our analyses, suggestively, associated with gene family expansion (i.e. a non-equilibrium SSDL process Figure S21). However with the taxon-sampling employed here, we find that gene count data alone do not provide enough signal to distinguish a flexible SSDL model from a DLWGD model. Posterior predictive model simulations indicate a similarly fair fit as for the dicots data set (Figure S22), with 32 and 29 out of 39 summary statistics within the 95% central mass of the marginal posterior predictive distribution for the analysis with *δ* = 0.05 and *δ* = 0.1 respectively.

## Discussion

In this paper we continued our previous work on the statistical inference of ancient WGD events from comparative genomic data (Zwaenepoel and Van de Peer 2019). Building on the initial idea of Rabier, Ta, and Ané (2013), we model gene family evolution by small-scale duplication and loss (SSDL) using a phylogenetic linear birth-death process and exploit the statistical deviation from such a process to infer ancient WGDs. In contrast to the approaches taken in these previous studies, we here do not assume a particular WGD configuration, and infer the number and locations of WGDs in a species tree from the data using reversible jump MCMC under a flexible hierarchical model of gene family evolution. In our previous work, we showed that using simple birth-death models for the SSDL process, in particular models assuming constant rates of duplication and loss across the entire phylogeny, seriously compromises reliable inference of ancient WGDs. Simulations showed that false positive rates for WGD inference become unacceptably high when the constant rates assumption is violated (Zwaenepoel and Van de Peer 2019), which we deem likely to be the rule rather than the exception for most phylogenies. On the other hand, employing more complicated, hierarchical models with branch-wise duplication and loss rates may compromise our ability to detect WGDs from gene count or gene tree information, with the assumed SSDL process sufficiently flexible to capture a WGD signature. In the Bayesian inference scheme this tradeoff translates to a certain degree of prior sensitivity, with assumptions on the rate evolution process potentially affecting our posterior inferences.

In this work we are confronted with the very same challenges. Using a flexible model for the SSDL process is vital to prevent false positive WGD inferences, but concomitantly results in a statistical approach with limited power to detect true WGD events for long branches or WGD events with small associated retention rates. In our multivariate models of duplication and loss rate evolution this flexibility is embodied by the covariance matrix of the branch-wise duplication and loss rates, which affects both the amount of rate heterogeneity across branches and the difference between the duplication and loss rate for any particular branch. We further optionally constrain the latter by using an additional prior on the expected number of lineages per ancestral lineage under the SSDL process, which has the advantage of being more intelligible than the prior covariance parameter, therefore allowing easier application of informative priors. Nevertheless, even when using fairly informative priors in the latter form we find that information from gene count data often does not provide decisive support for putative WGD events previously described in the literature.

These statistical problems of power and prior sensitivity are in the first place due to the kind of data employed for tackling the problem of WGD inference. Indeed, the same issues are relevant in some form or another whenever inference is based on phylogenomic data – whether in the form of gene trees of multi-copy gene families (as in Zwaenepoel and Van de Peer (2019), Li et al. (2018), 1KP initiative (2019) etc.) or gene counts (as in Rabier, Ta, and Ané (2013) and this study) – and ignores information from genome structure. By using a rich model we seek to explicitly account for the most relevant sources of variation in this data when assessing the history of WGDs in a phylogeny, and we are able to assess the effects of particular assumptions of the SSDL process on our WGD inferences. We believe the Bayesian approach advocated here compells us to embrace the inherent uncertainty of our inferences based on this kind of data, while at the same time allowing WGD inference in a coherent and biologically meaningful framework. The usage of gene trees instead of gene counts may further increase the power for WGD detection (Zwaenepoel and Van de Peer 2019), and the accuracy of duplication and loss rate estimates. However, this comes at the expense of a less efficient algorithm for computing the likelihood and a much more involved and computationally intensive data preparation phase (which in the Whale approach of Zwaenepoel and Van de Peer (2019) involves obtaining a sample from the posterior distribution of gene tree topologies for every gene family). Using gene counts instead could allow us to employ broader sets of taxa, which may be beneficial, as improved taxon sampling could result in a higher power to detect ancient WGDs, although this has to be studied in more detail. To us, the most fruitful avenue for future research appears to involve an integration of information from genome structure in the probabilistic phylogenomic framework we adopt here; leveraging the obvious differences between the SSDL and WGD process at the level of gene synteny or co-linearity. Lastly, we note that while posterior predictive simulations suggest that the DLWGD model with branch-wise rates provides a reasonable fit to the data, there may be room for improvement on the modeling side as well. However, more complicated models (e.g. involving non-linear BDPs (Crawford, Minin, and Suchard 2014) or rate heterogeneity across families) will generally be associated with even higher computational demands. Another potentially interesting improvement in terms of modeling would be to account for incompletely sampled genomes and assembly or annotation errors (as for instance in Han et al. (2013) or Rabier, Ta, and Ané (2013)) in the Bayesian approach we adopt here.

Computational problems are however more of an issue than the statistical intricacies discussed above, which are after all reflecting the very evolutionary processes of interest. Whereas we have implemented the likelihood evaluation routines and rjMCMC algorithm in an (to the best of our knowledge) efficient manner and exploit distributed computing architectures to accelerate computation of the likelihood across families; we nevertheless had to restrict our analyses to relatively limited sets of taxa and subsets of the full data. Furthermore, whereas mixing within a particular WGD configuration tends to be very efficient irrespective of the data set size, larger data sets tend to be associated with poor mixing across model space in the rjMCMC, with the acceptance probability for trans-dimensional moves dropping rapidly with increasing data set size. This confers a risk that for larger data sets an incomplete view of model uncertainty is obtained, and it forces us to sample longer chains to ensure decent mixing of the chain across model space; making the problem even more computationally intractable. Instead of investing computer power in the analysis of more gene families, it may be worthwhile to instead focus on adding more taxa, as this could similarly improve the statistical power for WGD inference while not affecting mixing efficiency. We note that as a follow-up analysis, a resulting set of putative WGDs could further be tested extensively using a fixed dimension MCMC sampler in a manner analogous to Rabier, Ta, and Ané (2013) and Zwaenepoel and Van de Peer (2019), possibly with a larger data set. This could result in a more accurate approximation of the posterior distribution of the duplication, loss and retention rates under a particular DLWGD model compared to the approximated distribution from the rjMCMC sample. Finally, we have in this study only considered WGD, whereas at least some ancestral polyploidization events are associated with higher multiplication levels such as the *γ* triplication in eudicots (Jiao et al. 2012) and the Solanaceae triplication (Tomato Genome Consortium 2012). It is possible to extend the DLWGD model to higher multiplication levels (Rabier, Ta, and Ané 2013), and in the Bayesian approach we adopt here, the multiplication level could in principle be included as an additional parameter, or alternatively, reversible jump moves for different multiplication levels could be included. We did not however explore this additional layer of complexity in our current study and defer this to future work.

In summary, we present a fully model-based approach for WGD detection in a phylogenetic context using a flexible hierarchical model of gene family evolution. Posterior inference is based on trans-dimensional MCMC using the reversible-jump Metropolis-Hastings algorithm. We model variation of duplication and loss rates using a multivariate approach, and we note that this in principle would allow to model correlated evolution with other quantitative traits as in Lartillot and Poujol (2010), although we did not explore this in our study and defer this to future research efforts. Through simulations and analyses of comparative genomic data from flowering plants we investigated the reliability and power of our new approach, and provide further insights in the statistical inference problem of detecting ancient WGDs from phylogenomic data. We find that when a flexible model of gene family evolution is assumed, the power for WGD inference is rather limited, with strong support across all analyses only for those WGDs with a very strong signature (for instance the WGDs in the *C. quinoa, P. trichocarpa, A. thaliana, M. acuminata* and *E. guineensis* lineages in our analyses). For some other putative WGDs (*M. truncatula*, *S. lycopersicum* and *Z. mays*) we find that our inferences are sensitive to prior assumptions, indicating that some caution is warranted when applying the proposed methods. In general, we believe performing multiple analyses under progressively more restrictive priors may provide insights in which WGD hypotheses are supported under which assumptions on the background SSDL process. Computationally, our rjMCMC approach is challenging, especially for large data sets, and the Bayesian inference machinery underlying this work is an obvious target for future improvements. Eventually, we hope that the methods presented here enable statistically better informed inferences of ancient WGDs, and can contribute to an improved understanding of this key and increasingly appreciated process in genome evolution.

## Acknowledgements

Arthur Zwaenepoel is supported by a PhD fellowship of the Research Foundation Flanders (FWO).

## Supplementary material

**Figure S1:**
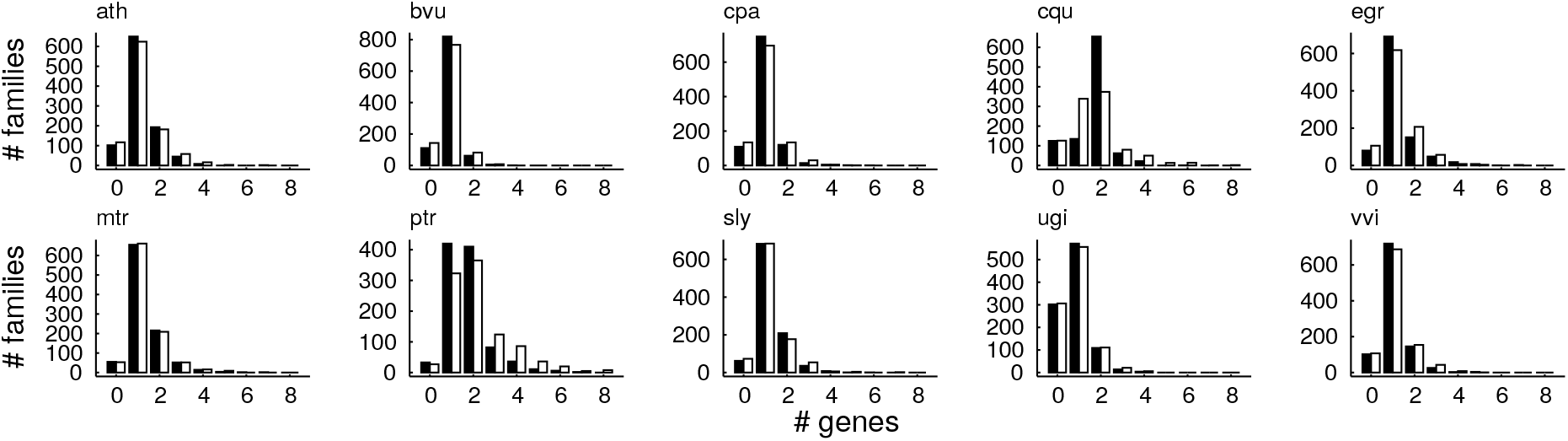
Comparison of the distribution of the number of genes of each species in a gene family for the observed data (1000 gene families across 10 plant species) (in black) with simulations from the posterior predictive distribution (in white) under the BM model of duplication and loss rate variation. Considering for example *Chenopodium quinoa* (cqu) or *P. trichocarpa* (ptr) (two species with a well-described highly preserved ancient WGD), and comparing for instance with *Vitis vinifera* (vvi) or *Carica papaya* (cpa) (two species with no known WGDs in their recent evolutionary past), these simulations clearly illustrate that WGDs are a major source of model violation when modeling gene family evolution by a flexible linear birth-death process.

**Figure S2:**
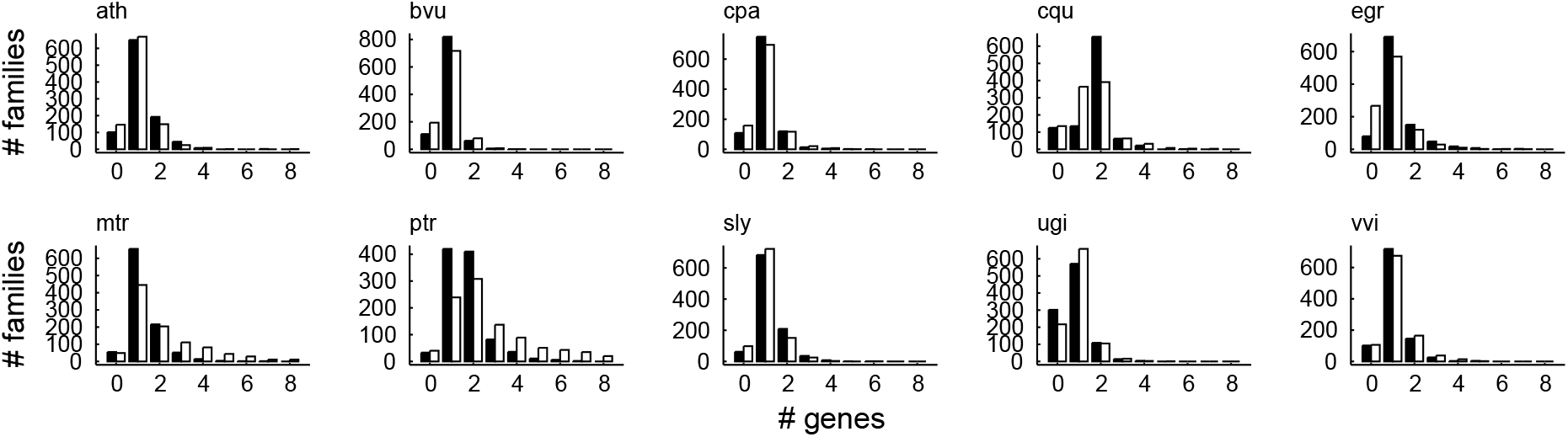
As in Figure S1 but under the IR model for rate variation across the species tree

**Figure S3:**
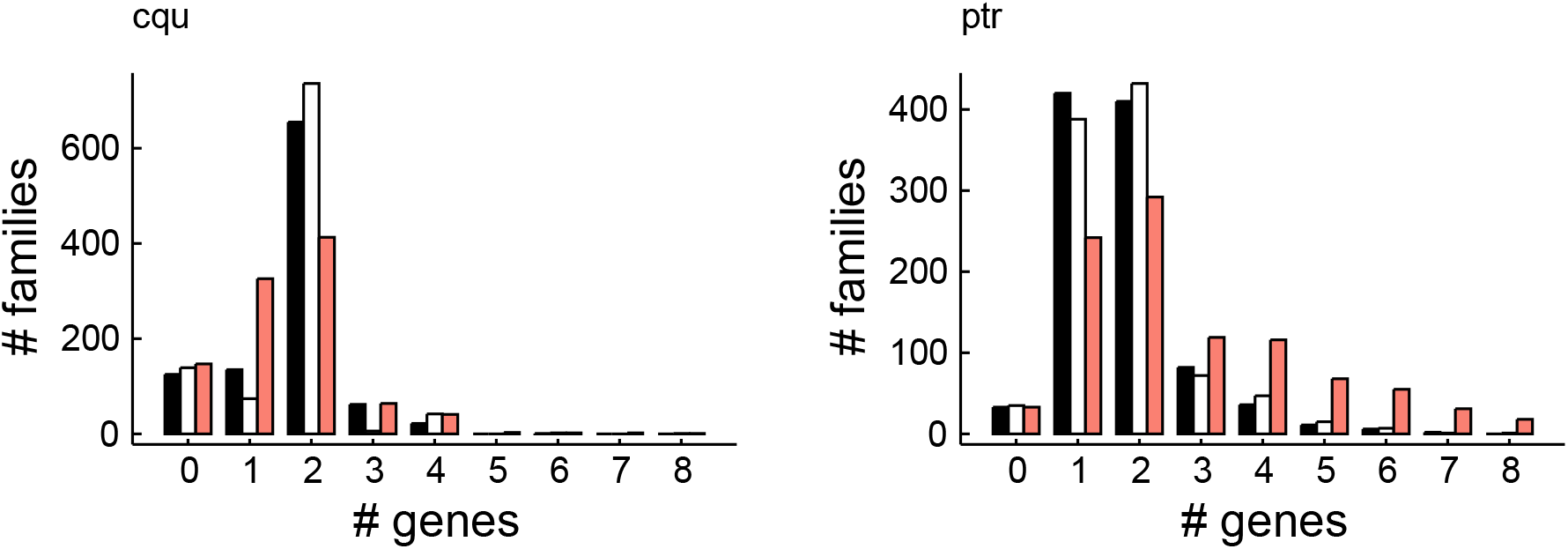
As in Figure S2 but now the SSDL-only model is shown in red and we include in white posterior predictive simulations under a model including a WGD on the branches leading to *C.quinoa* (cqu) and *P. trichocarpa* (ptr).

**Figure S4:**
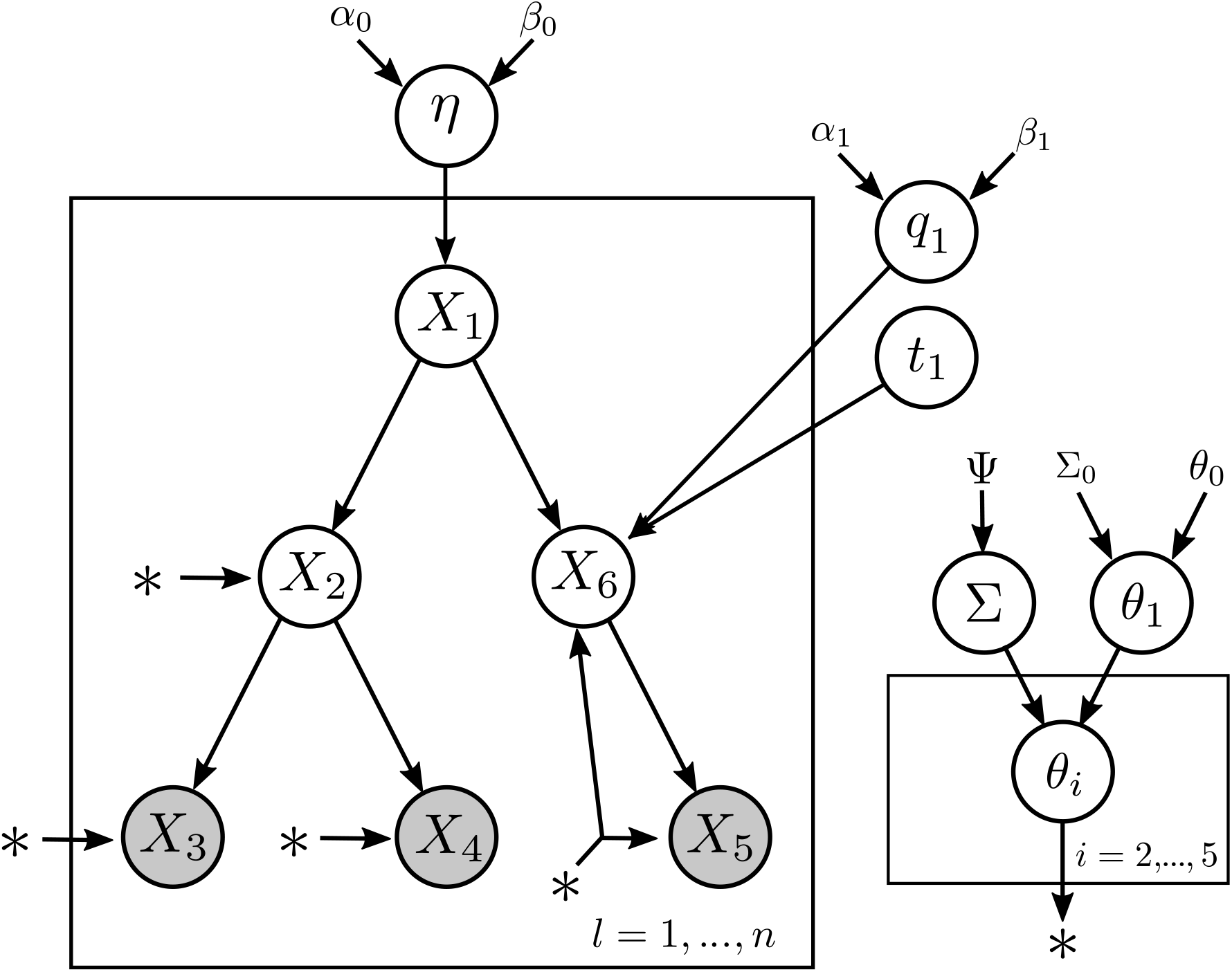
Probabilistic graphical model (PGM) illustrating the hierarchical structure of the IR-model for a hypothetical three-taxon species tree with one WGD node (*X*_6_).

**Figure S5:**
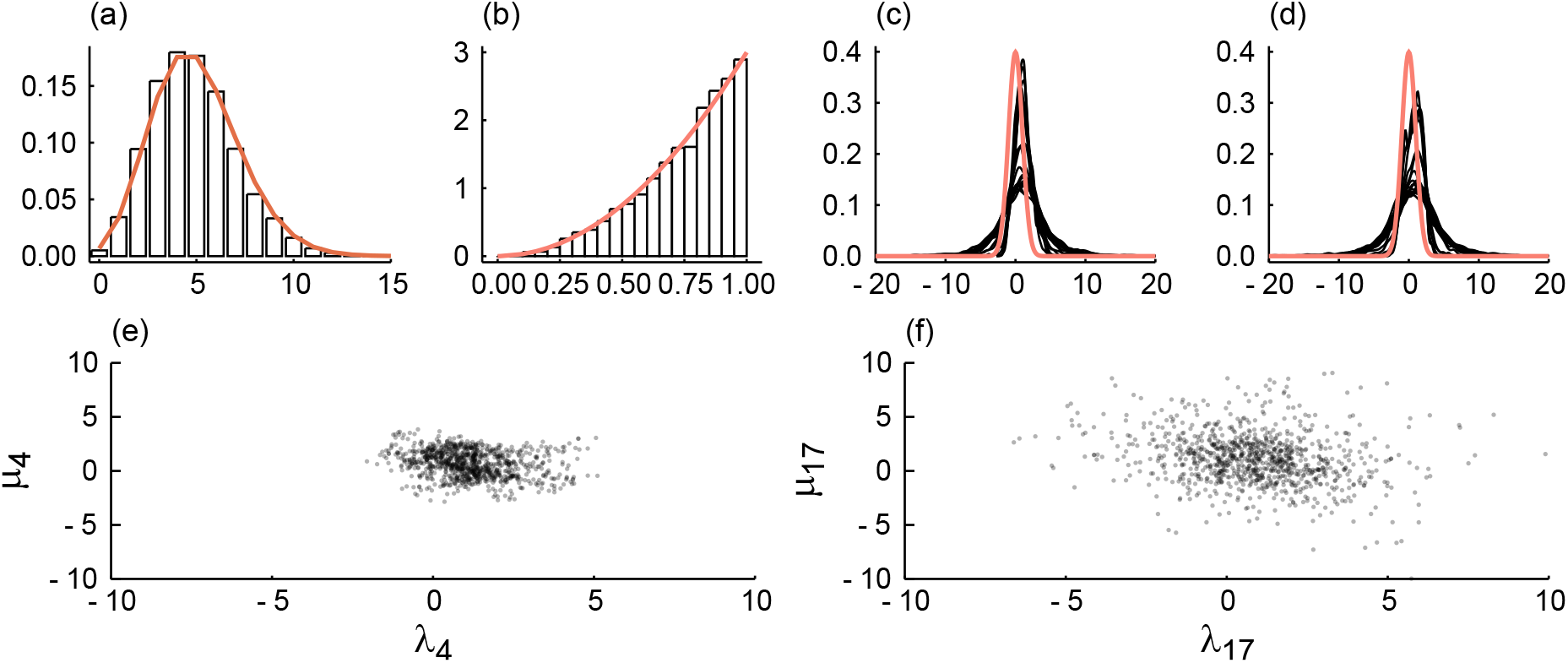
MCMC samples from the BM prior. (a) The number of WGDs *k*, with a Poisson prior (in orange), (b) *η* with a Beta prior (in orange), (c) duplication rates, with in orange the prior on the root duplication rate log *λ*_1_, (d) as in (c) but for loss rates (e) scatter plot of log *μ*_4_ vs. log *λ*_4_ and (f) as in (e) but for node 17.

**Figure S6:**
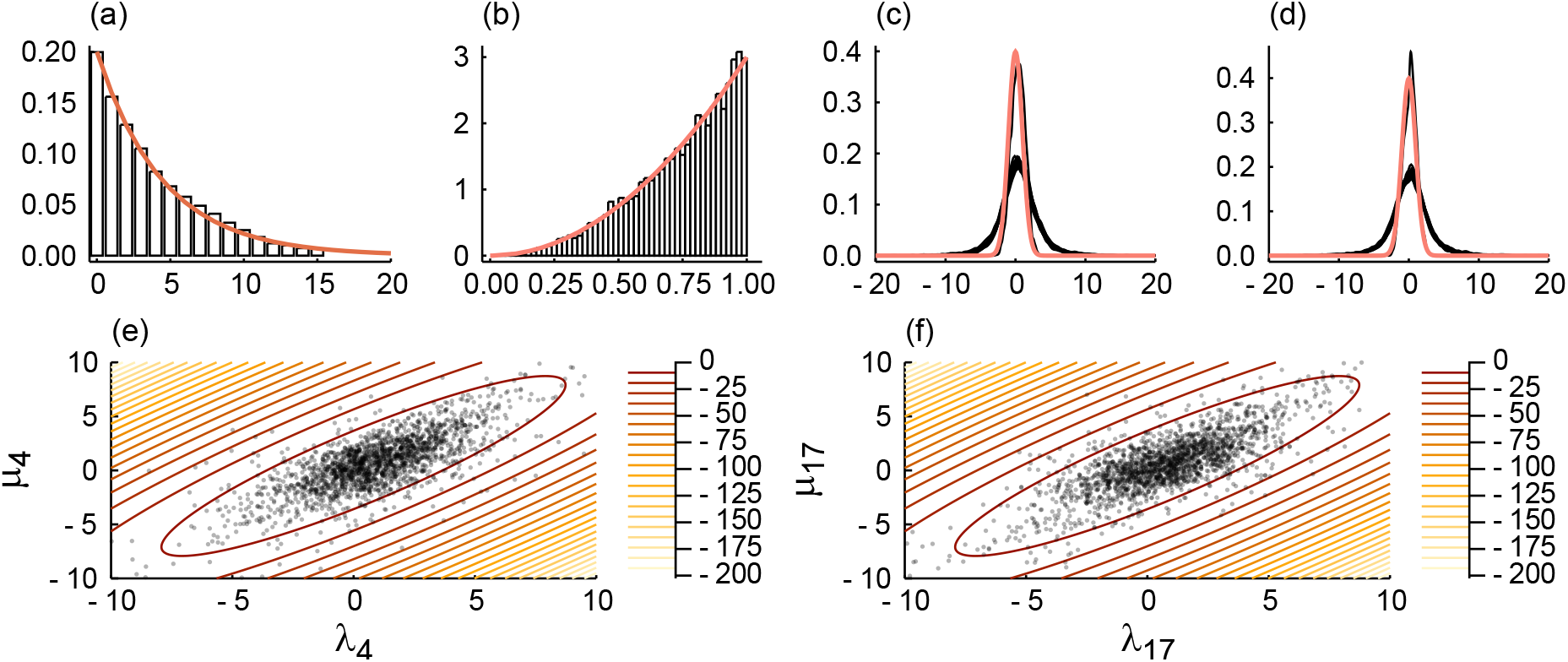
MCMC samples from the IR prior with a Geometric(0.25) prior with upper bound of 15 on the number of WGDs, and a prior covariance matrix with non-zero non-diagonal entries. See Figure S5 for explanation of the panels. The overlayed contour plot in panels (e) and (f) shows the log density Multivariate Normal distribution with mean *θ*_0_ and covariance matrix Σ_0_ = Ψ.

**Figure S7:**
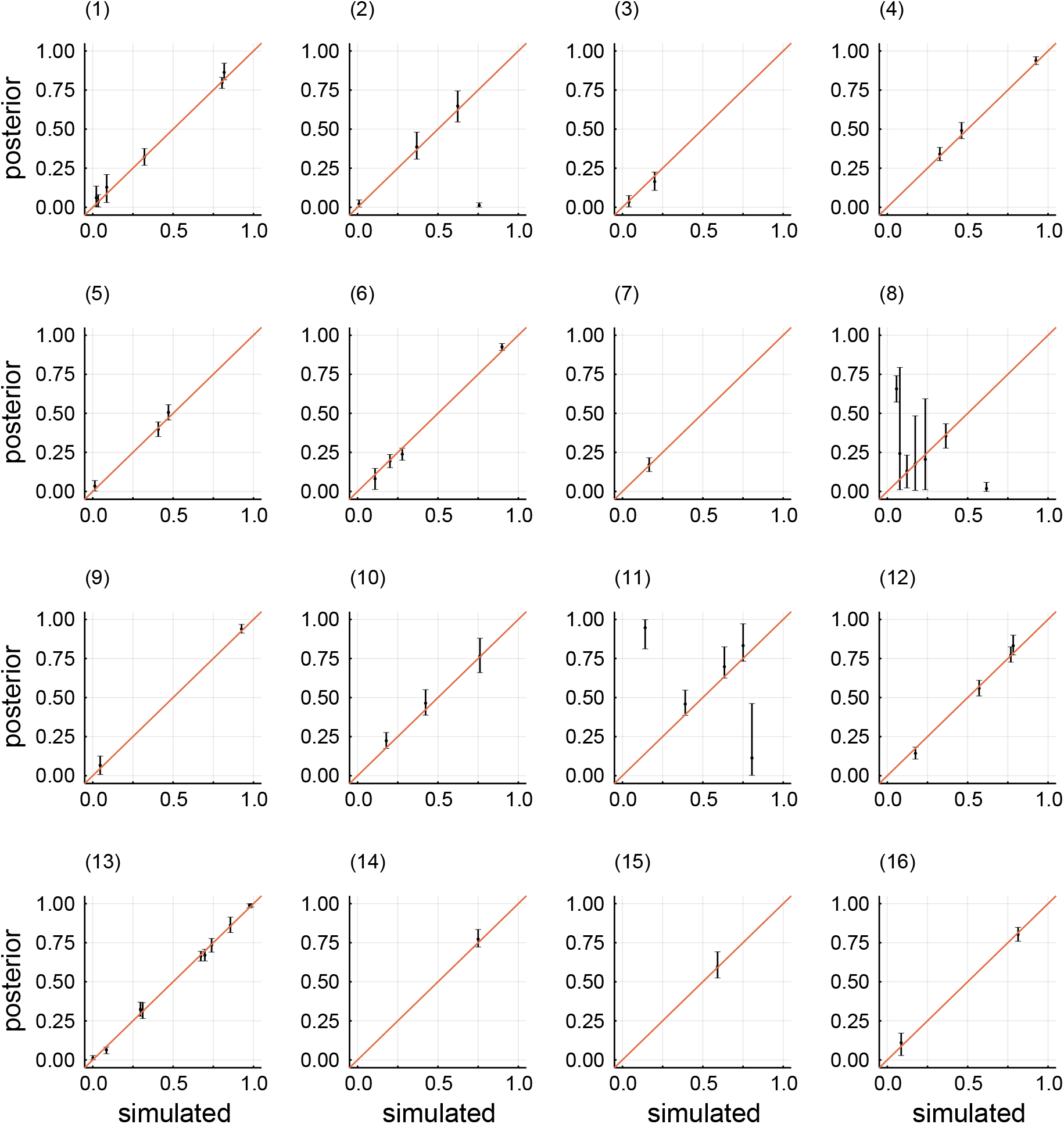
Posterior mean estimates and associated 95% credibility intervals for retention rates, as in Figure 1 but here we show results for each of the 16 replicates (that included WGDs) separately. We note that replicate 8 and 11 show the issue where retention rates for WGDs on the same branch are swapped (see main text).

**Figure S8:**
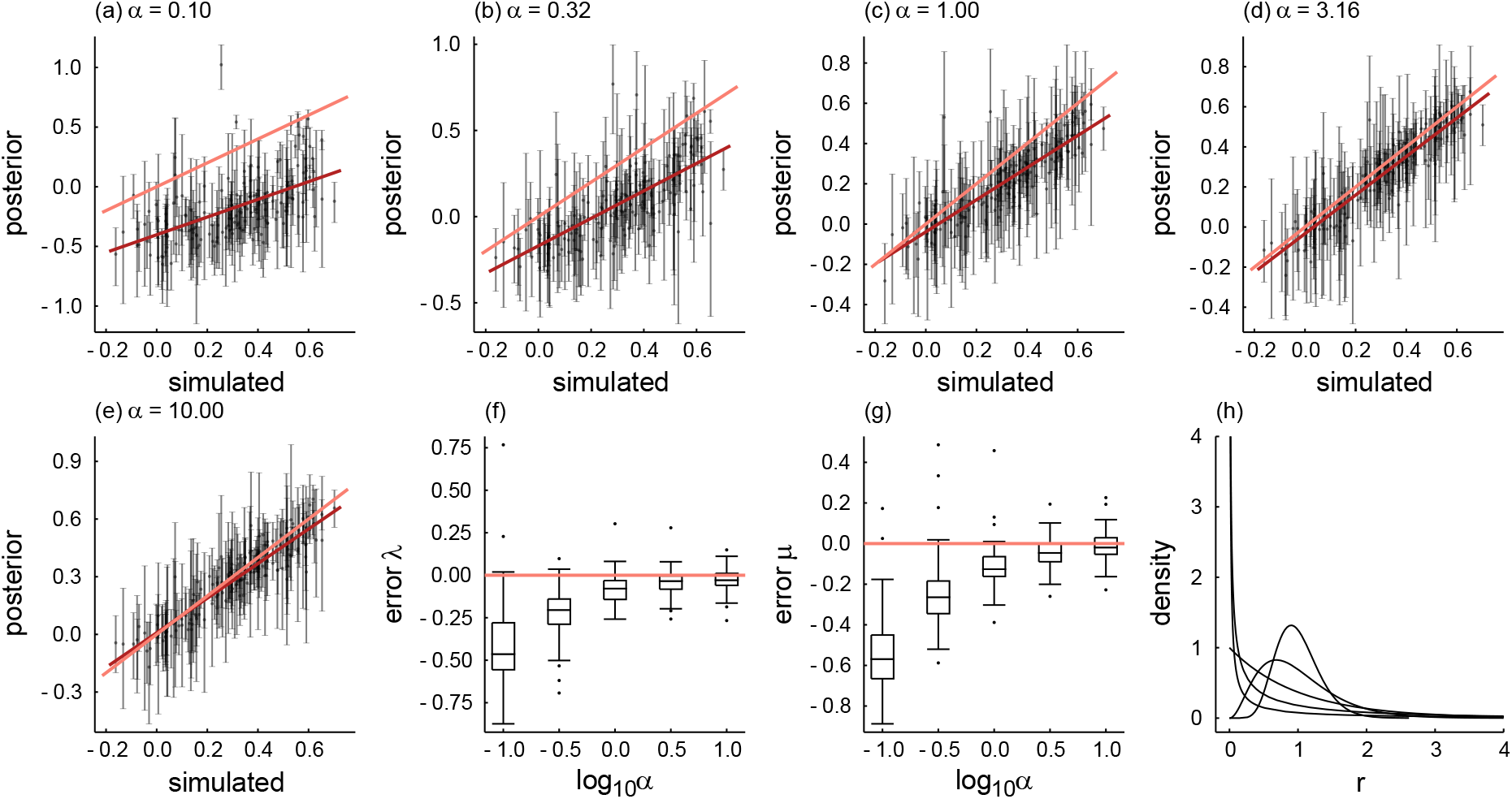
The effects of family × lineage variation on the posterior inference of duplication and loss rates. Simulations with Gamma distributed family-specific relative rates and without WGDs were performed for different shapes of the Gamma distribution, i.e. Gamma(*α*, 1/*α*) (such that the mean is 1) for different *α*. Posterior inference was performed using the fixed-dimensional MCMC sampler, assuming constant rates of duplication and loss across families (but not across lineages). Duplication and loss rate estimates (on a log_10_ scale) for five replicate simulations of *N* = 500 families are shown pooled together in panels (a-e). Panels (f) and (g) show the distributions of the difference between the posterior geometric mean and true value. Panel (h) illustrates the different shapes of the Gamma distribution considered in these simulations

**Figure S9:**
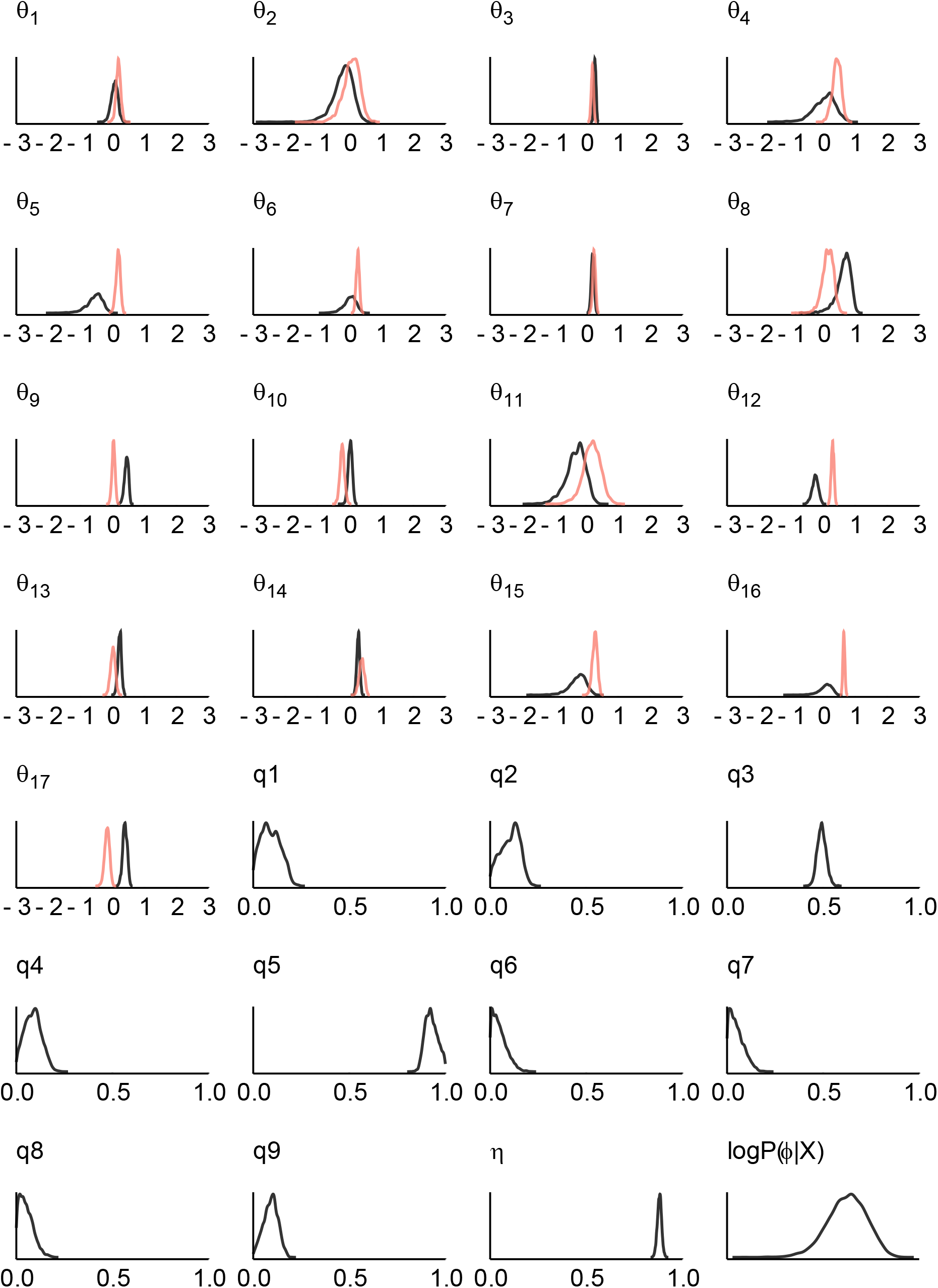
Approximate marginal posterior densities for branch-wise duplication and loss rates (*θ_i_*, black and red respectively, on a log_10_ scale), retention rates *q_j_* and *η* for the dicot data set obtained with a fixed-dimensional MCMC sampler for a nine-WGD model. Duplication and loss rates for the second set of simulations shown in Figure 2 were drawn from the joint posterior distribution associated with these results.

**Figure S10:**
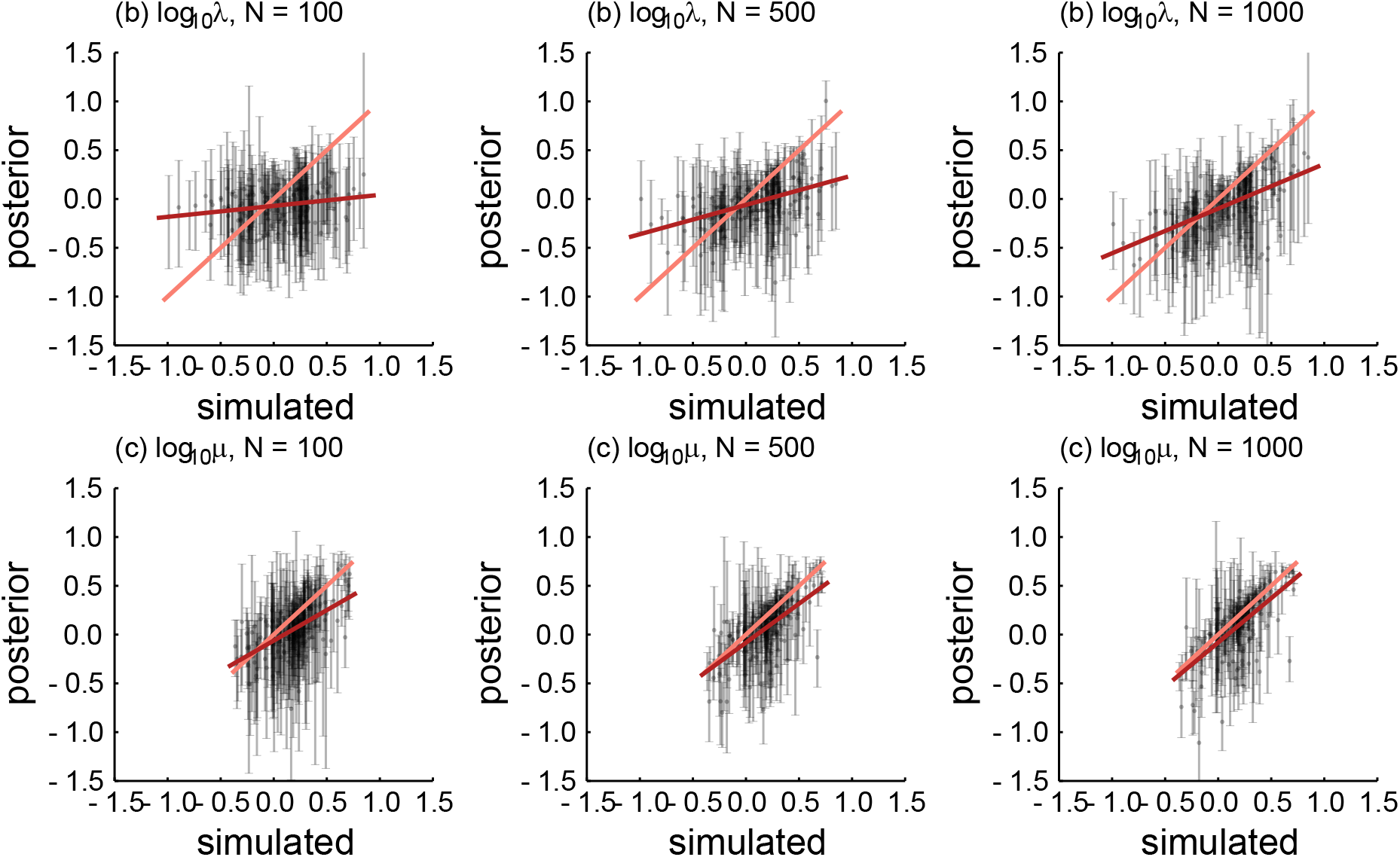
Posterior mean duplication and loss rates with 95% credibility intervals for the second set of simulations (i.e. the data sets simulated from the approximate joint posterior for the nine dicots data set acquired with the fixed-dimensional sampler), obtained using the rjMCMC sampler with prior covariance matrix Ψ = 0.2 *I*_2_. In orange the true relationship is shown whereas in red the least squares regression of posterior means on the true simulated values is shown. Results for all branch-wise rates for 10 simulated replicates are shown pooled together. Note how the variation in the duplication rates is much higher than than the variation in the loss rates.

**Figure S11:**
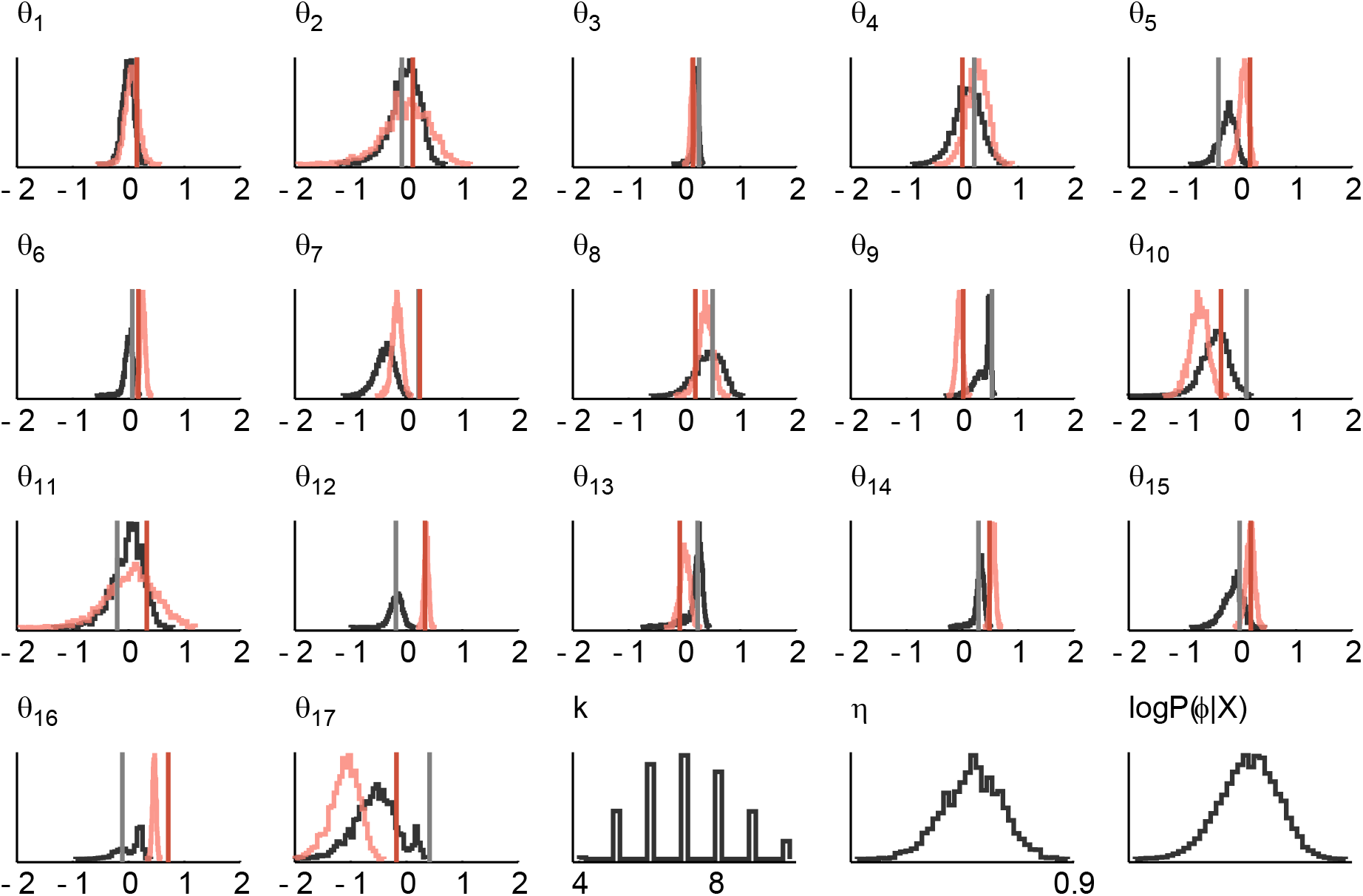
Approximate marginal posterior densities for one of the simulated replicates (*N* = 1000, Ψ = 0.2 *I*_2_, where simulated duplication and loss rates were drawn from the joint posterior for the dicots data set under a fixed model, see main text for more details) obtained with the rjMCMC algorithm. The branch-wise duplication and loss rates (*θ_i_*, on a log_10_ scale) are shown in black and red respectively. See also Figure S12.

**Figure S12:**
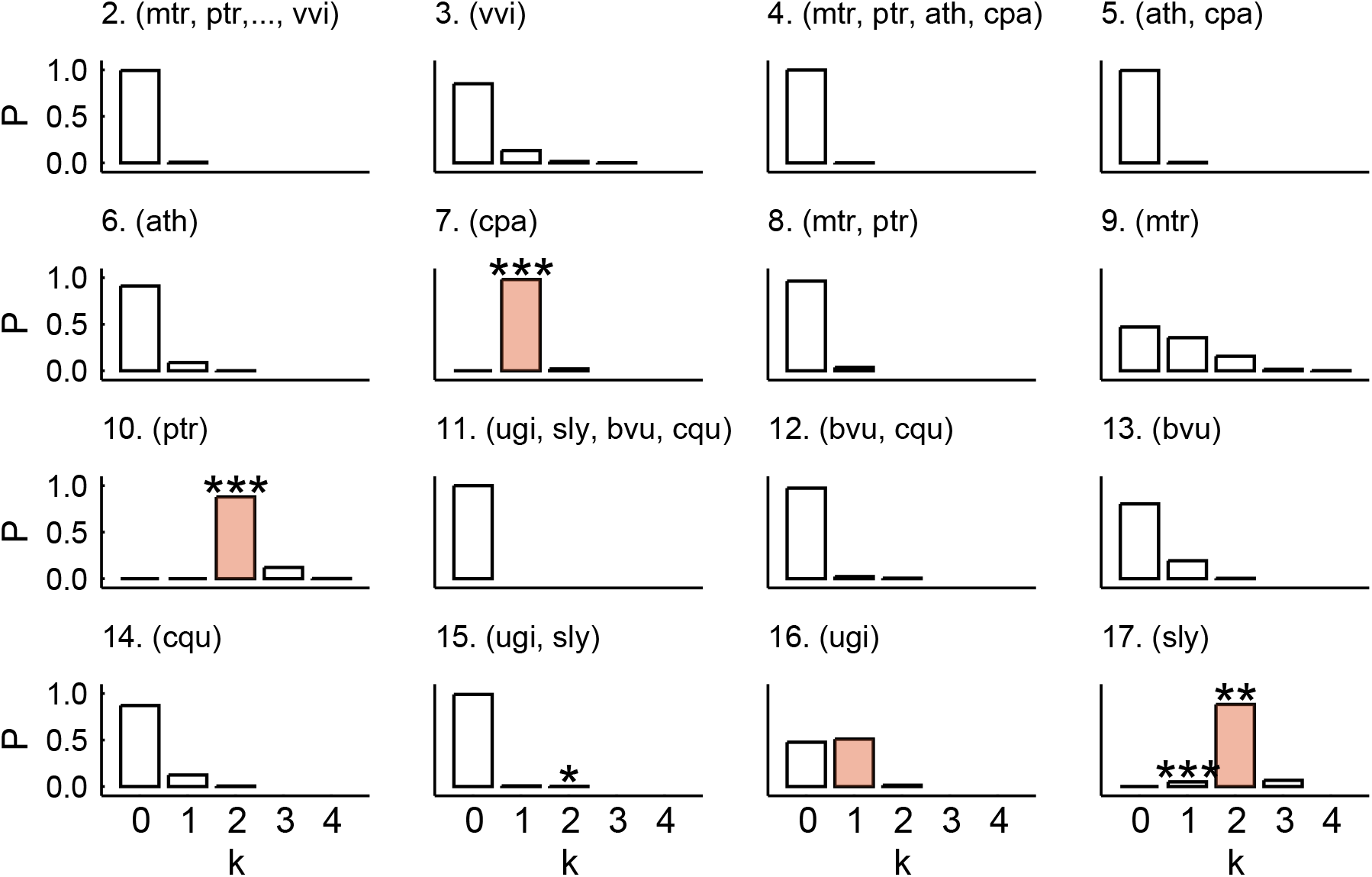
Posterior probabilities for the number of WGDs on each branch for the same simulated data as in Figure S11. Asterisks indicate the magnitude of the Bayes factor (*) 0.5 < log_10_ *K* < 1; (**) 1 < log_10_ *K* < 2; (***) log_10_ *K* > 2. The true number of WGDs are marked in red.

**Figure S13:**
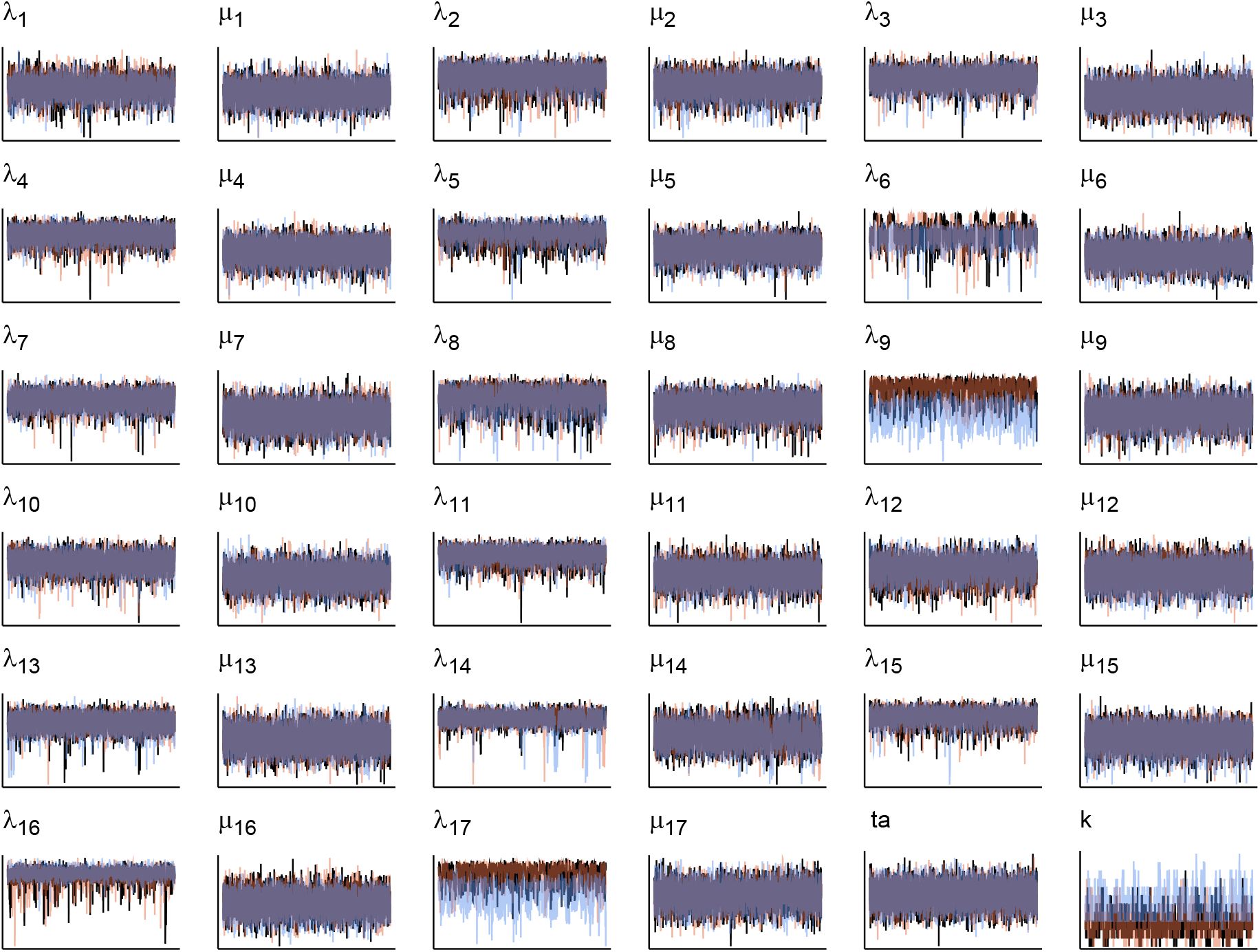
Trace plots for two realizations of the rjMCMC chain with *δ* = 0.1 (black and red) and one realization with *δ* = 0.05 (blue) based on 1000 gene families for the nine dicot species tree. Log-transformed duplication (*λ*) and loss (*μ*) rates are shown for each branch of the species tree. *k* denotes the number of WGDs in the model. Traces are based on 50000 iterates from the rjMCMC algorithm.

**Figure S14:**
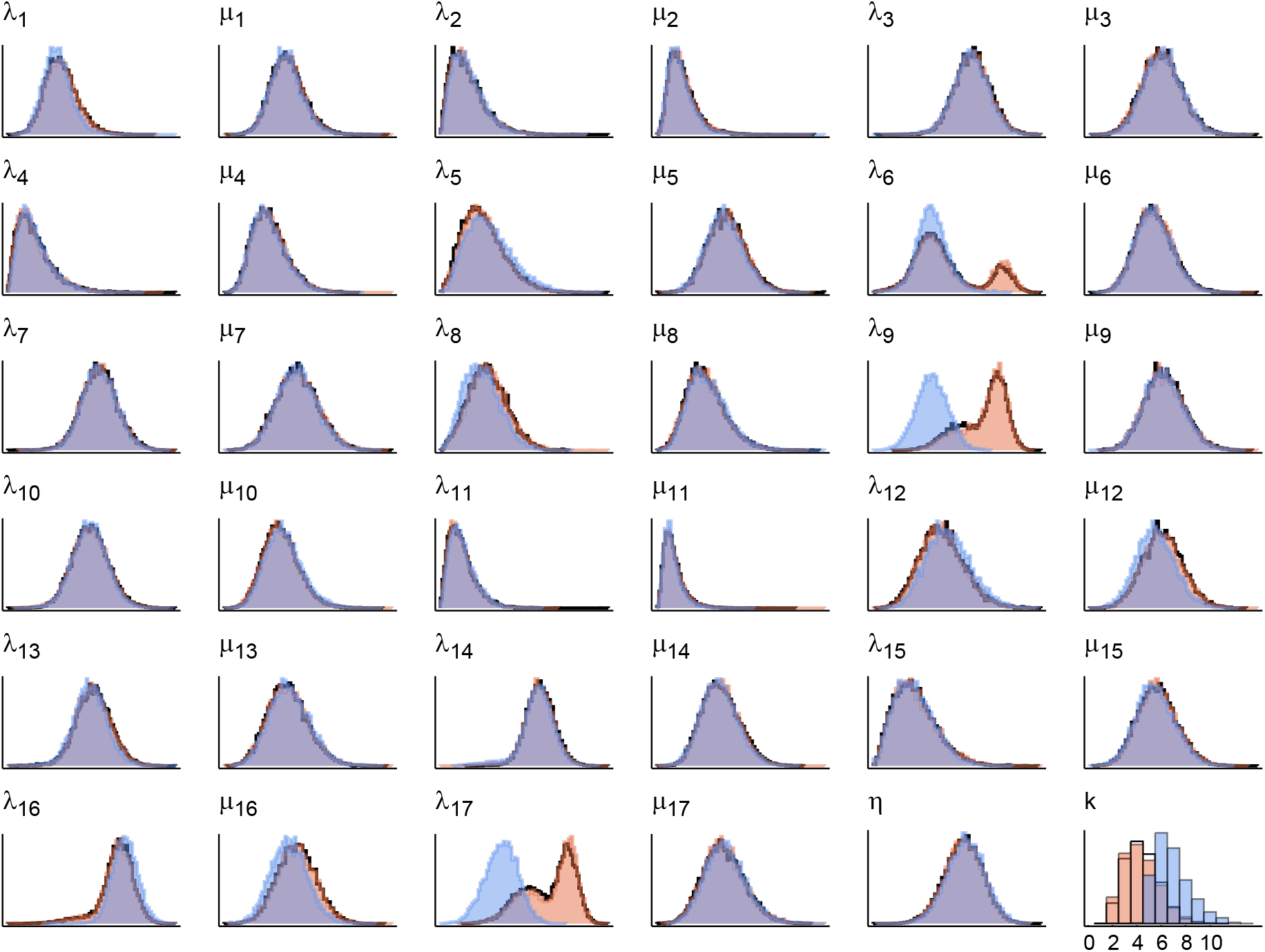
Approximate marginal posterior distributions corresponding to the trace plots shown in Figure S13.

**Figure S15:**
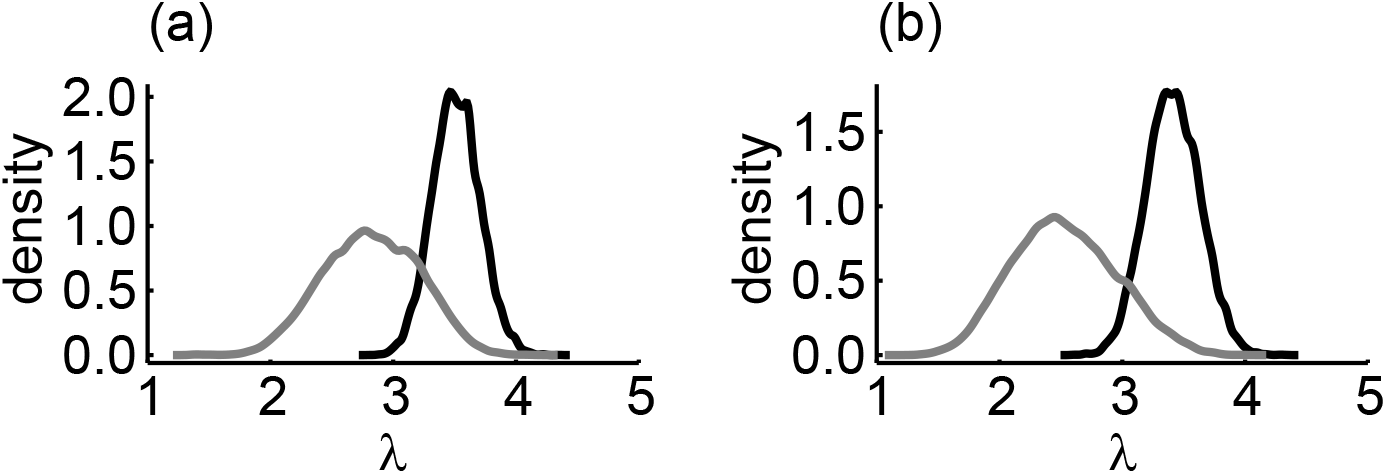
Conditional posterior duplication rates for (a) *Medicago* and (b) *Solanum*, conditioning on the no-WGD model (black) and one-WGD model (gray) for the results using the prior with *δ* = 0.1. The *x*-axis is on a scale of number of events per gene lineage per billion years.

**Figure S16:**
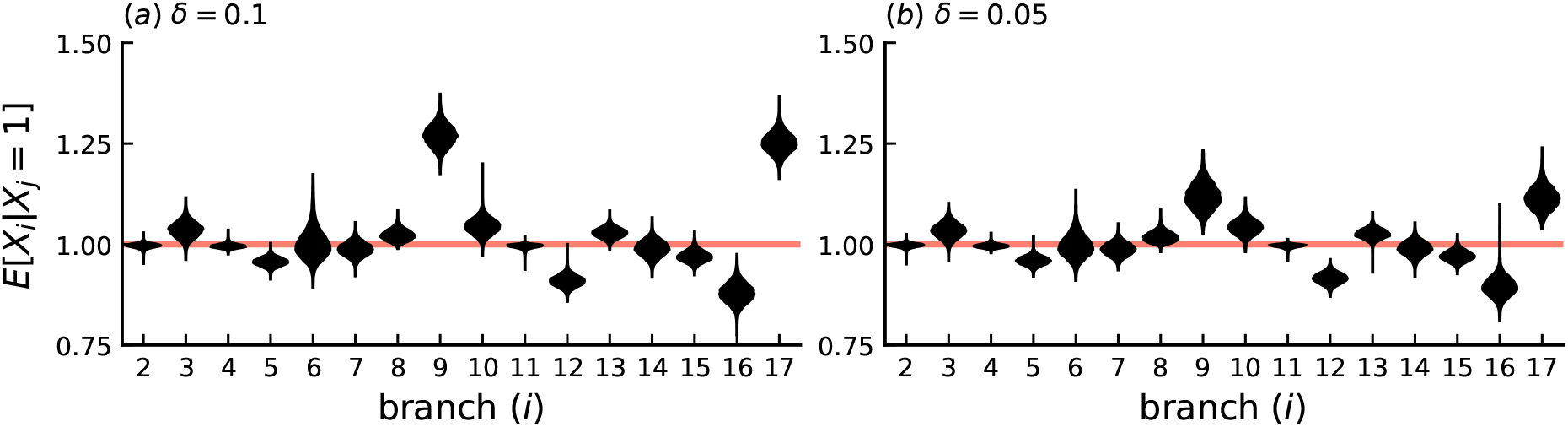
Posterior distributions for the expected number of genes per ancestral gene at the end of each branch under the SSDL process alone for the dicot dataset, showing results for two different prior settings, (a) *δ* = 0.1 and (b) *δ* = 0.05. For correspondence of branch numbers to clades, refer to Figure 3. The marginal posterior is shown over all different WGD models in the rjMCMC sample.

**Figure S17:**
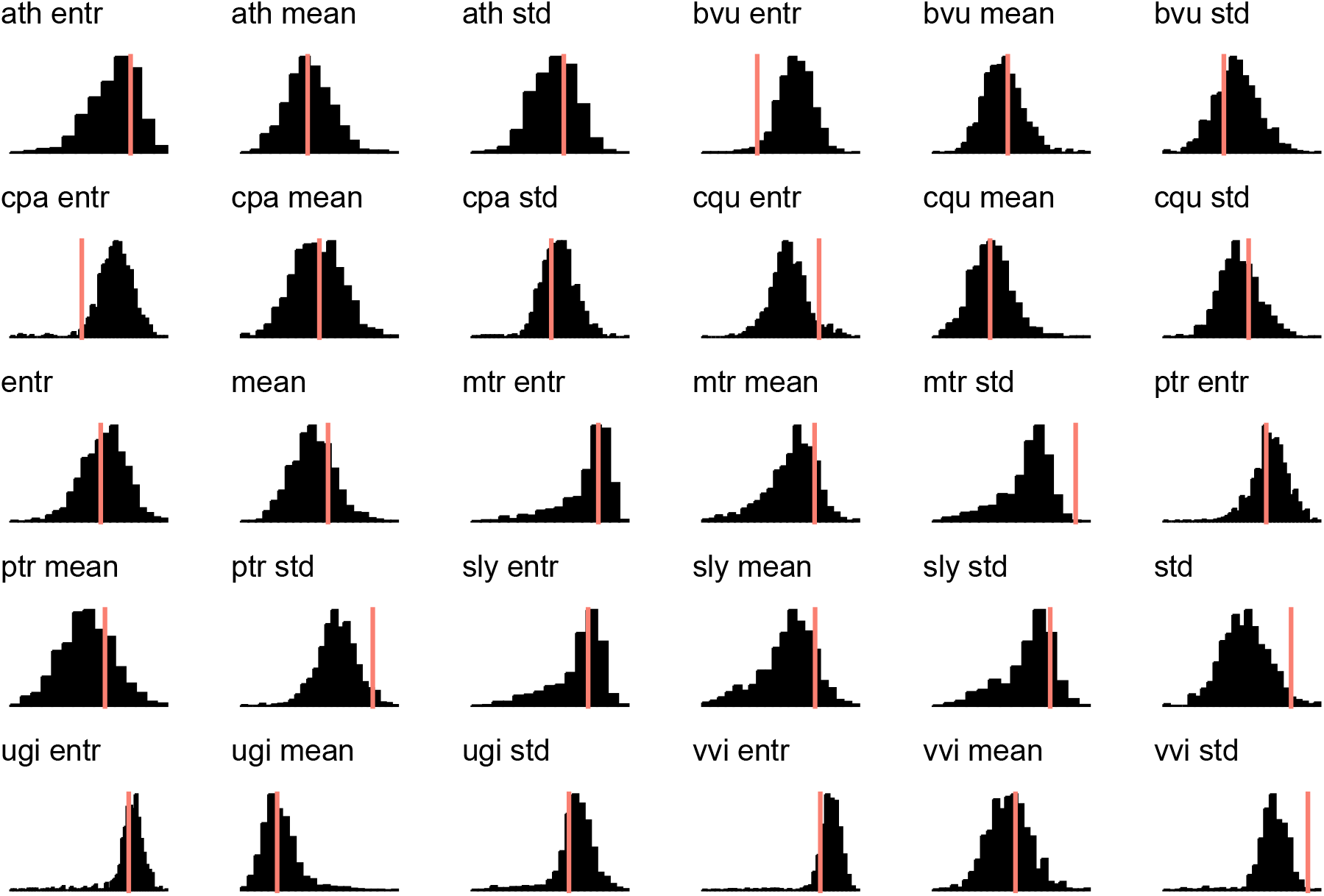
Posterior predictive distributions (histograms) and observed values (vertical lines) for 30 summary statistics of the phylogenetic profile matrix for the dicots data set with the *δ* = 0.1 prior. Summary statistics are the mean, standard deviation (std) and entropy (entr) of the number of genes across families in the leaves of the species tree, as well as the total number of genes in a family.

**Figure S18:**
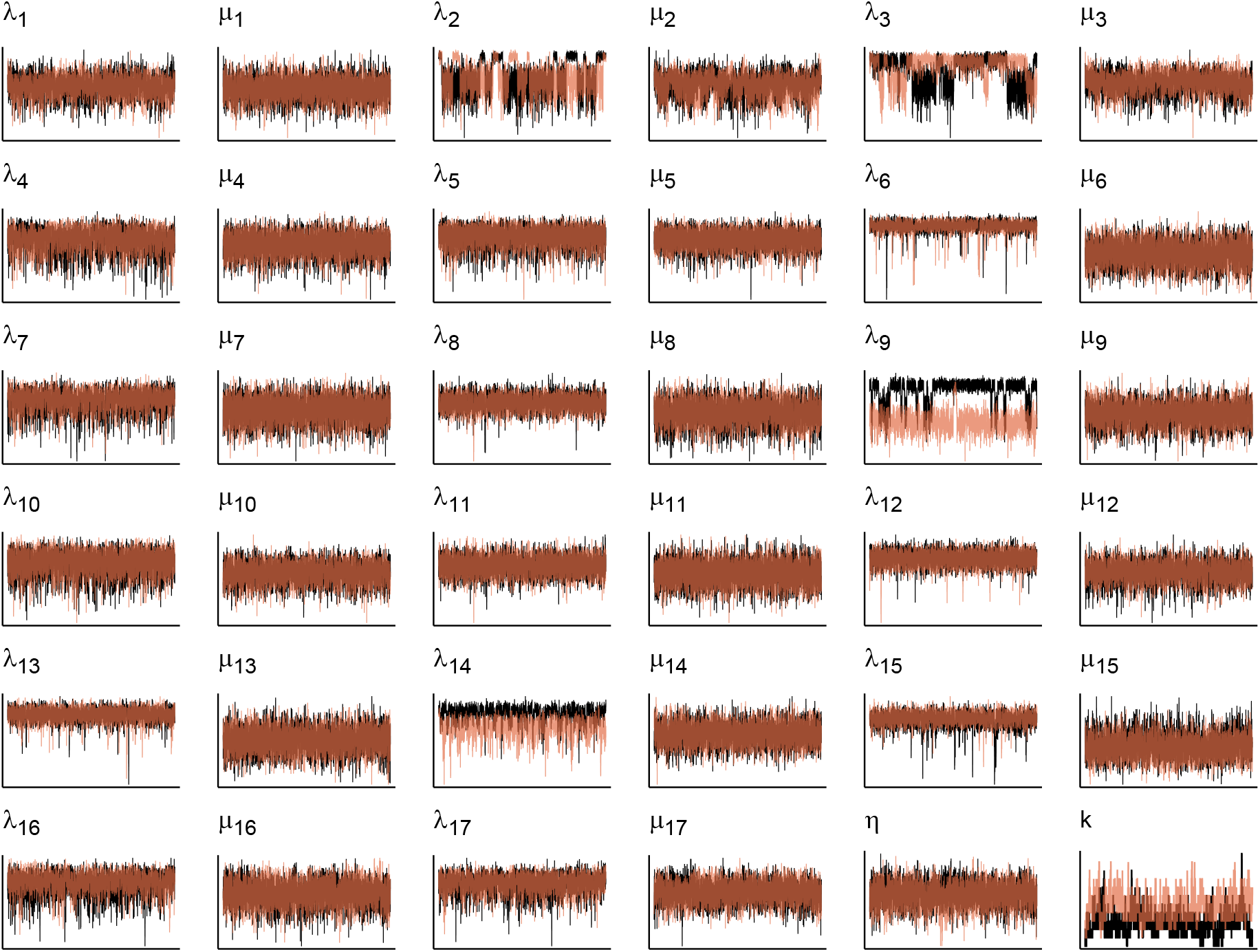
Trace plots for two realizations of the rjMCMC chain with 1000 gene families for the monocot species tree. In black a chain with *δ* = 0.1 is shown whereas in orange a chain with *δ* = 0.05 is shown. Log-transformed duplication (*λ*) and loss (*μ*) rates are shown for each branch of the species tree. *k* denotes the number of WGDs in the model. Traces are based on 50000 iterates from the rjMCMC algorithm.

**Figure S19:**
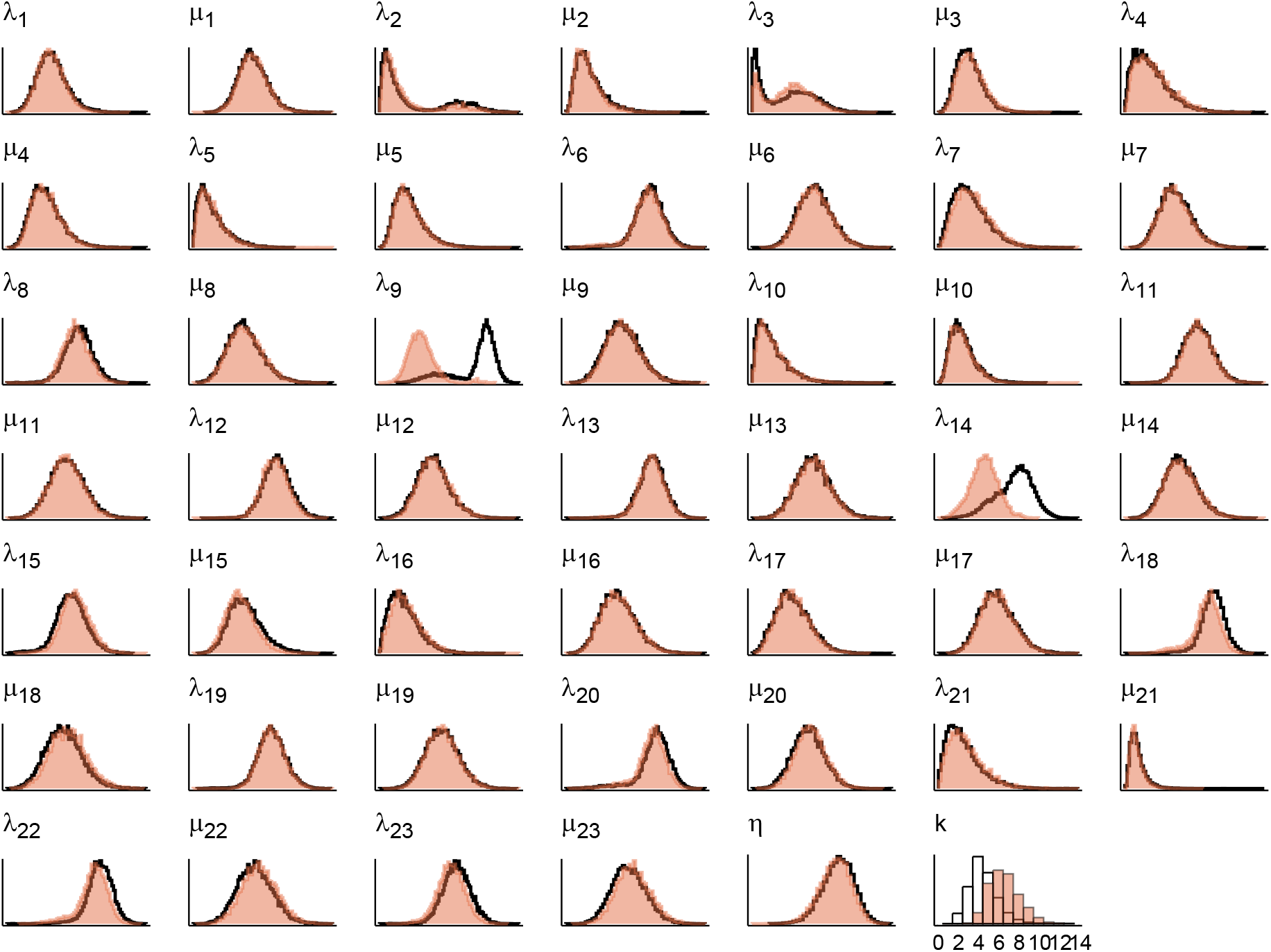
Approximate marginal posterior distributions for two realizations (black and red histograms) of the rjMCMC chain with 1000 gene families for the monocot species tree. Interpretation is as in Figure S18.

**Figure S20:**
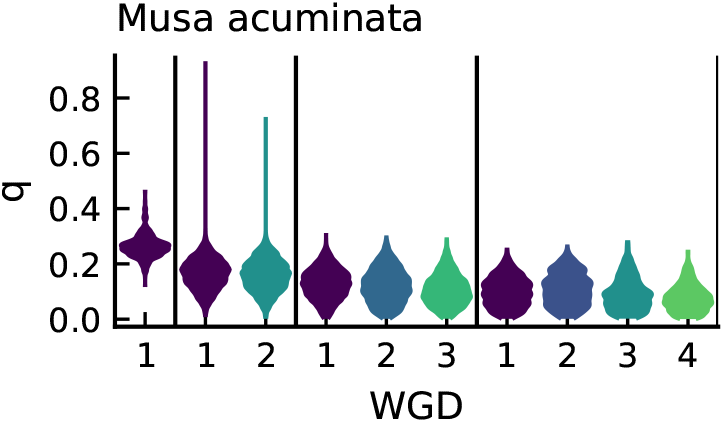
Marginal posterior distributions of retention rates for the *Musa acuminata* WGDs across different models. Vertical lines separate different models with different numbers of WGDs.

**Figure S21:**
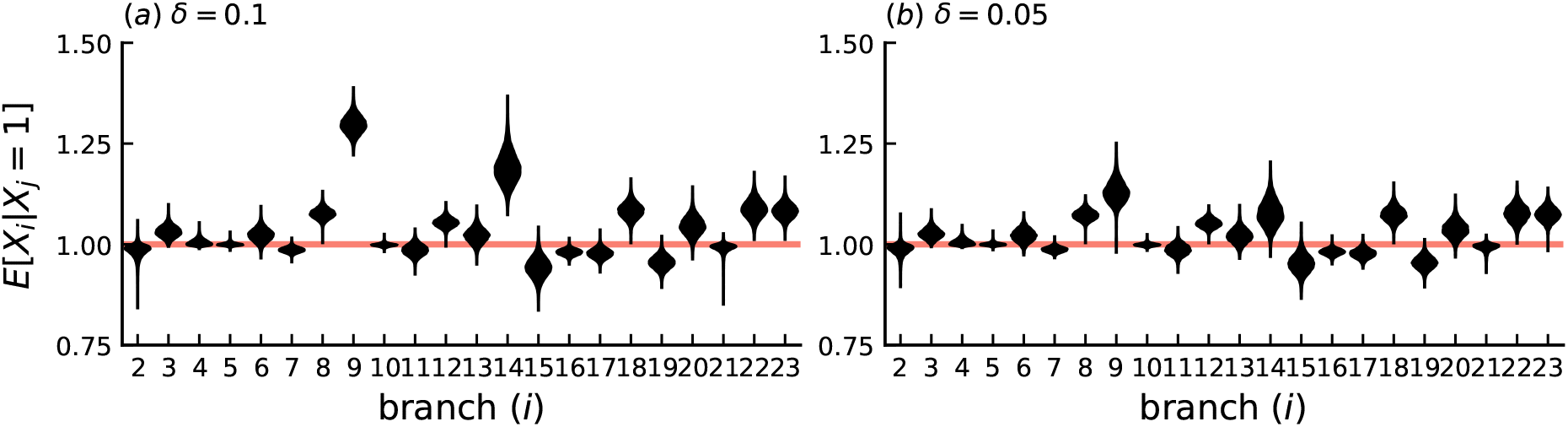
Posterior distributions for the expected number of genes per ancestral gene at the end of each branch under the SSDL process alone for the monocot dataset, showing results for two different prior settings, (a) *δ* = 0.1 and (b) *δ* = 0.05. For correspondence of branch numbers to clades, refer to Figure 4, 5. The marginal posterior is shown over all different WGD models in the rjMCMC sample.

**Figure S22:**
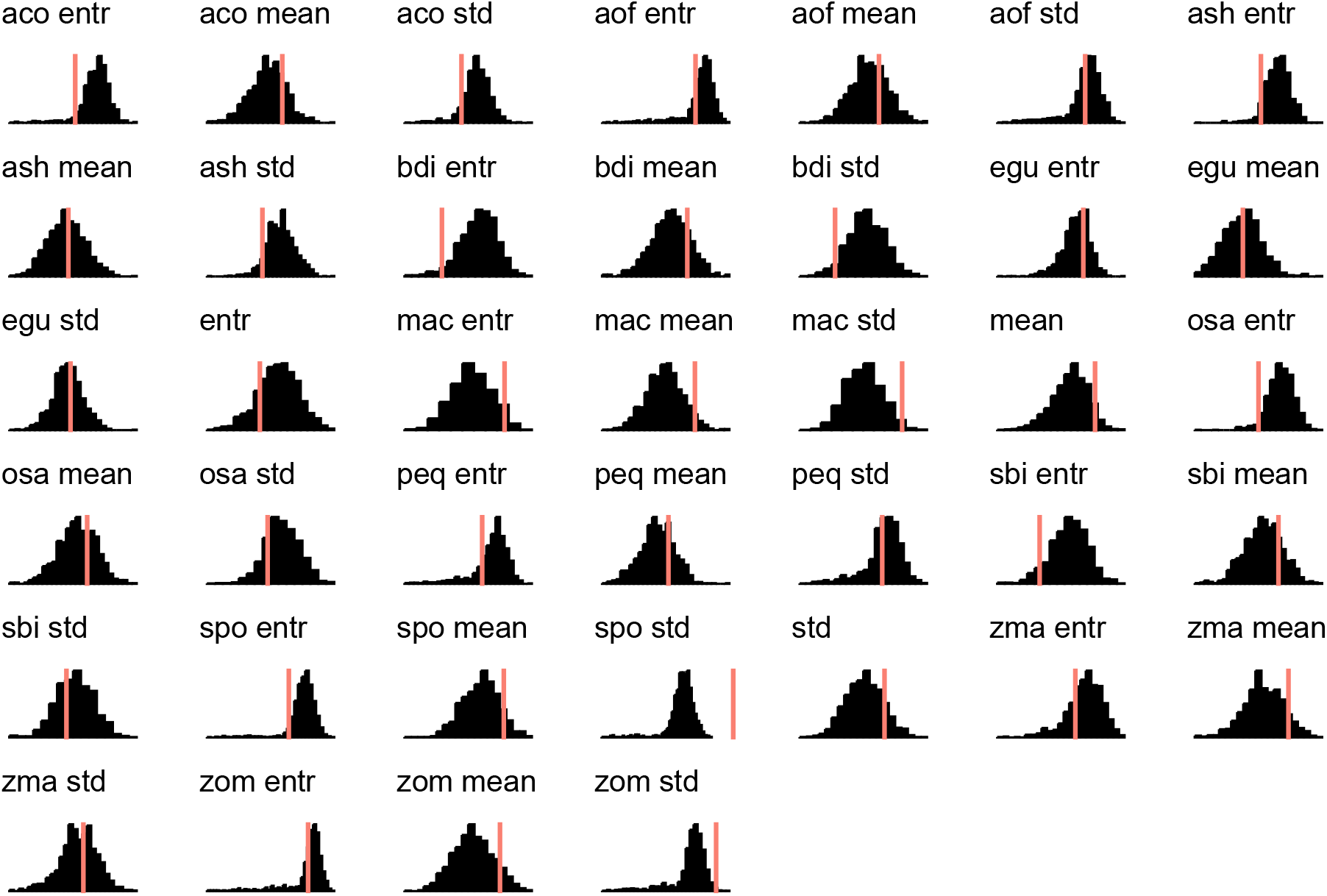
Posterior predictive distributions (histograms) and observed values (vertical lines) for 39 summary statistics of the phylogenetic profile matrix for the monocots data set. Summary statistics are the mean, standard deviation (std) and entropy (entr) of the number of genes across families in the leaves of the species tree, as well as the total number of genes in a family. These results are based on the chain with *δ* = 0.05.

## References

1KP initiative. 2019. “One Thousand Plant Transcriptomes and Phylogenomics of Green Plants.” Nature 574: 679–85.

Bezanson, Jeff, Alan Edelman, Stefan Karpinski, and Viral B Shah. 2017. “Julia: A Fresh Approach to Numerical Computing.” SIAM Review 59 (1): 65–98.

Brooks, Stephen P, Paolo Giudici, and Gareth O Roberts. 2003. “Efficient Construction of Reversible Jump Markov Chain Monte Carlo Proposal Distributions.” Journal of the Royal Statistical Society: Series B (Statistical Methodology) 65 (1): 3–39.

Cai, Jing, Xin Liu, Kevin Vanneste, Sebastian Proost, Wen-Chieh Tsai, Ke-Wei Liu, Li-Jun Chen, et al. 2015. “The Genome Sequence of the Orchid Phalaenopsis Equestris.” Nature Genetics 47 (1): 65.

Carretero-Paulet, Lorenzo, Pablo Librado, Tien-Hao Chang, Enrique Ibarra-Laclette, Luis Herrera-Estrella, Julio Rozas, and Victor A Albert. 2015. “High Gene Family Turnover Rates and Gene Space Adaptation in the Compact Genome of the Carnivorous Plant Utricularia Gibba.” Molecular Biology and Evolution 32 (5): 1284–95.

Crawford, Forrest W, Vladimir N Minin, and Marc A Suchard. 2014. “Estimation for General Birth-Death Processes.” Journal of the American Statistical Association 109 (506): 730–47.

Csűrös, Miklós, and István Miklós. 2009. “Streamlining and Large Ancestral Genomes in Archaea Inferred with a Phylogenetic Birth-and-Death Model.” Molecular Biology and Evolution 26 (9): 2087–95.

D’Hont, Angélique, France Denoeud, Jean-Marc Aury, Franc-Christophe Baurens, Françoise Carreel, Olivier Garsmeur, Benjamin Noel, et al. 2012. “The Banana (Musa Acuminata) Genome and the Evolution of Monocotyledonous Plants.” Nature 488 (7410): 213.

Feller, William. 2015. “Die Grundlagen Der Volterraschen Theorie Des Kampfes Ums Dasein in Wahrscheinlichkeitstheoretischer Behandlung.” In Selected Papers I, 441–70. Springer.

Foster, Charles SP, Hervé Sauquet, Marlien Van der Merwe, Hannah McPherson, Maurizio Rossetto, and Simon YW Ho. 2016. “Evaluating the Impact of Genomic Data and Priors on Bayesian Estimates of the Angiosperm Evolutionary Timescale.” Systematic Biology 66 (3): 338–51.

Gelman, Andrew, John B Carlin, Hal S Stern, David B Dunson, Aki Vehtari, and Donald B Rubin. 2013. Bayesian Data Analysis. Chapman; Hall/CRC.

Green, Peter J. 1995. “Reversible Jump Markov Chain Monte Carlo Computation and Bayesian Model Determination.” Biometrika 82 (4): 711–32.

Hahn, Matthew W, Tijl De Bie, Jason E Stajich, Chi Nguyen, and Nello Cristianini. 2005. “Estimating the Tempo and Mode of Gene Family Evolution from Comparative Genomic Data.” Genome Research 15 (8): 1153–60.

Han, Mira V, Gregg WC Thomas, Jose Lugo-Martinez, and Matthew W Hahn. 2013. “Estimating Gene Gain and Loss Rates in the Presence of Error in Genome Assembly and Annotation Using Cafe 3.” Molecular Biology and Evolution 30 (8): 1987–97.

Harkess, Alex, Jinsong Zhou, Chunyan Xu, John E Bowers, Ron Van der Hulst, Saravanaraj Ayyampalayam, Francesco Mercati, et al. 2017. “The Asparagus Genome Sheds Light on the Origin and Evolution of a Young Y Chromosome.” Nature Communications 8 (1): 1279.

Huelsenbeck, John P, Bret Larget, and David Swofford. 2000. “A Compound Poisson Process for Relaxing the Molecular Clock.” Genetics 154 (4): 1879–92.

Ibarra-Laclette, Enrique, Eric Lyons, Gustavo Hernández-Guzmán, Claudia Anahí Pérez-Torres, Lorenzo Carretero-Paulet, Tien-Hao Chang, Tianying Lan, et al. 2013. “Architecture and Evolution of a Minute Plant Genome.” Nature 498 (7452): 94.

Jiao, Yuannian, Jim Leebens-Mack, Saravanaraj Ayyampalayam, John E Bowers, Michael R McKain, Joel McNeal, Megan Rolf, et al. 2012. “A Genome Triplication Associated with Early Diversification of the Core Eudicots.” Genome Biology 13 (1): R3.

Jiao, Yuannian, Jingping Li, Haibao Tang, and Andrew H Paterson. 2014. “Integrated Syntenic and Phylogenomic Analyses Reveal an Ancient Genome Duplication in Monocots.” The Plant Cell 26 (7): 2792–2802.

Kumar, Sudhir, Glen Stecher, Michael Suleski, and S Blair Hedges. 2017. “TimeTree: A Resource for Timelines, Timetrees, and Divergence Times.” Molecular Biology and Evolution 34(7): 1812–9.

Lartillot, Nicolas, and Raphaël Poujol. 2010. “A Phylogenetic Model for Investigating Correlated Evolution of Substitution Rates and Continuous Phenotypic Characters.” Molecular Biology and Evolution 28 (1): 729–44.

Li, Zheng, George P Tiley, Sally R Galuska, Chris R Reardon, Thomas I Kidder, Rebecca J Rundell, and Michael S Barker. 2018. “Multiple Large-Scale Gene and Genome Duplications During the Evolution of Hexapods.” Proceedings of the National Academy of Sciences 115 (18): 4713–8.

Li, Zheng, George P Tiley, Rebecca J Rundell, and Michael S Barker. 2019. “Reply to Nakatani and Mclysaght: Analyzing Deep Duplication Events.” Proceedings of the National Academy of Sciences 116(6): 1819–20.

Librado, Pablo, Filipe G Vieira, and Julio Rozas. 2011. “BadiRate: Estimating Family Turnover Rates by Likelihood-Based Methods.” Bioinformatics 28 (2): 279–81.

Liu, Liang, Lili Yu, Venugopal Kalavacharla, and Zhanji Liu. 2011. “A Bayesian Model for Gene Family Evolution.” BMC Bioinformatics 12 (1): 426.

Long, Manyuan, Nicholas W VanKuren, Sidi Chen, and Maria D Vibranovski. 2013. “New Gene Evolution: Little Did We Know.” Annual Review of Genetics 47: 307–33.

Lynch, Michael. 2007. The Origins of Genome Architecture. Vol. 98. Sinauer Associates Sunderland, MA.

Ming, Ray, Robert VanBuren, Ching Man Wai, Haibao Tang, Michael C Schatz, John E Bowers, Eric Lyons, et al. 2015. “The Pineapple Genome and the Evolution of Cam Photosynthesis.” Nature Genetics 47 (12): 1435.

Muller, Hermann J. 1936. “Bar Duplication.” Science 83 (2161): 528–30.

Nakatani, Yoichiro, and Aoife McLysaght. 2019. “Macrosynteny Analysis Shows the Absence of Ancient Whole-Genome Duplication in Lepidopteran Insects.” Proceedings of the National Academy of Sciences 116(6): 1816–8.

Novozhilov, Artem S, Georgy P Karev, and Eugene V Koonin. 2006. “Biological Applications of the Theory of Birth-and-Death Processes.” Briefings in Bioinformatics 7 (1): 70–85.

Olsen, Jeanine L, Pierre Rouzé, Bram Verhelst, Yao-Cheng Lin, Till Bayer, Jonas Collen, Emanuela Dattolo, et al. 2016. “The Genome of the Seagrass Zostera Marina Reveals Angiosperm Adaptation to the Sea.” Nature 530 (7590): 331.

Rabier, Charles-Elie, Tram Ta, and Cécile Ané. 2013. “Detecting and Locating Whole Genome Duplications on a Phylogeny: A Probabilistic Approach.” Molecular Biology and Evolution 31 (3): 750–62.

Rannala, Bruce, and Ziheng Yang. 2013. “Improved Reversible Jump Algorithms for Bayesian Species Delimitation.” Genetics 194 (1): 245–53.

Roberts, Gareth O, and Jeffrey S Rosenthal. 2009. “Examples of Adaptive Mcmc.” Journal of Computational and Graphical Statistics 18 (2): 349–67.

Ruiz-Orera, Jorge, Pol Verdaguer-Grau, José Luis Villanueva-Cañas, Xavier Messeguer, and M Mar Albà. 2018. “Translation of Neutrally Evolving Peptides Provides a Basis for de Novo Gene Evolution.” Nature Ecology & Evolution 2 (5): 890.

Singh, Rajinder, Meilina Ong-Abdullah, Eng-Ti Leslie Low, Mohamad Arif Abdul Manaf, Rozana Rosli, Rajanaidu Nookiah, Leslie Cheng-Li Ooi, et al. 2013. “Oil Palm Genome Sequence Reveals Divergence of Interfertile Species in Old and New Worlds.” Nature 500 (7462): 335.

Soltis, Pamela S, and Douglas E Soltis. 2016. “Ancient Wgd Events as Drivers of Key Innovations in Angiosperms.” Current Opinion in Plant Biology 30: 159–65.

Tasdighian, Setareh, Michiel Van Bel, Zhen Li, Yves Van de Peer, Lorenzo Carretero-Paulet, and Steven Maere. 2017. “Reciprocally Retained Genes in the Angiosperm Lineage Show the Hallmarks of Dosage Balance Sensitivity.” The Plant Cell 29 (11): 2766–85.

Tiley, George P, Cecile Ane, and J Gordon Burleigh. 2016. “Evaluating and Characterizing Ancient Whole-Genome Duplications in Plants with Gene Count Data.” Genome Biology and Evolution 8 (4): 1023–37.

Tomato Genome Consortium. 2012. “The Tomato Genome Sequence Provides Insights into Fleshy Fruit Evolution.” Nature 485 (7400): 635.

Van Bel, Michiel, Tim Diels, Emmelien Vancaester, Lukasz Kreft, Alexander Botzki, Yves Van de Peer, Frederik Coppens, and Klaas Vandepoele. 2017. “PLAZA 4.0: An Integrative Resource for Functional, Evolutionary and Comparative Plant Genomics.” Nucleic Acids Research 46 (D1): D1190–D1196.

Van de Peer, Yves, Eshchar Mizrachi, and Kathleen Marchal. 2017. “The Evolutionary Significance of Polyploidy.” Nature Reviews Genetics 18 (7): 411.

Wang, Wenkin, G Haberer, H Gundlach, C Gläßer, TCLM Nussbaumer, MC Luo, A Lomsadze, et al. 2014. “The Spirodela Polyrhiza Genome Reveals Insights into Its Neotenous Reduction Fast Growth and Aquatic Lifestyle.” Nature Communications 5: 3311.

Yang, Ziheng. 1994. “Maximum Likelihood Phylogenetic Estimation from Dna Sequences with Variable Rates over Sites: Approximate Methods.” Journal of Molecular Evolution 39 (3): 306–14.

Zhang, Guo-Qiang, Ke-Wei Liu, Zhen Li, Rolf Lohaus, Yu-Yun Hsiao, Shan-Ce Niu, Jie-Yu Wang, et al. 2017. “The Apostasia Genome and the Evolution of Orchids.” Nature 549 (7672): 379.

Zhang, Li, Yan Ren, Tao Yang, Guangwei Li, Jianhai Chen, Andrea R Gschwend, Yeisoo Yu, et al. 2019. “Rapid Evolution of Protein Diversity by de Novo Origination in Oryza.” Nature Ecology & Evolution 3 (4): 679.

Zwaenepoel, Arthur, Zhen Li, Rolf Lohaus, and Yves Van de Peer. 2019. “Finding Evidence for Whole Genome Duplications: A Reappraisal.” Molecular Plant 12 (2): 133–36.

Zwaenepoel, Arthur, and Yves Van de Peer. 2019. “Inference of Ancient Whole-Genome Duplications and the Evolution of Gene Duplication and Loss Rates.” Molecular Biology and Evolution 36 (7): 1384–1404.

